# Prioritization in working memory reduces interference via beta band-linked stimulus transformations

**DOI:** 10.64898/2026.02.24.707753

**Authors:** Jacqueline M. Fulvio, Bradley R. Postle

## Abstract

We studied the effects of prioritization in a two-step retrocuing task in which male and female humans hold two items in working memory, and the item not cued by the first cue cannot be dropped because it may be prioritized by the second cue. In Experiment 1, using a dense sampling procedure, recall performance oscillated at 15 Hz (“low-beta”) in the prioritization task, in comparison to 20 Hz (“higher-beta”) in a matched neutral-cue task. Furthermore, the prioritized item was shielded from bias exerted by the uncued item, as well as by items from the previous trial. In Experiment 2, we sought EEG correlates of these effects while participants performed variants of the two tasks. The prioritization cue uniquely triggered a phase reset at 15 Hz and an increase in oscillatory peaks at the “low-beta” frequency. Burst analysis ruled out bursting as a possible underlying factor. Self-correlation analysis of the EEG time series revealed that the prioritization cue triggered greater reorganization of the whole-topography EEG pattern than the neutral cue, and for the uncued item than for the cued item. These results provide evidence that the shielding effects of prioritization may arise from the transformation of the not-prioritized item into an “unprioritized” state that is implemented and maintained by a mechanism that cycles in the “low-beta” frequency range (centered on roughly 15 Hz).

**Significance statement:** When multiple items are held in working memory, the brain must prioritize the subset of information needed to guide impending action while also retaining the other information that remains potentially relevant for future actions. Through behavior and electroencephalography, we show that prioritizing one item in working memory shields it from interference from the not-prioritized item and that this shielding may be implemented by transforming the representational state of the unprioritized item. Behavioral signatures of prioritization and the underlying neural dynamics both cycle at a frequency in the low-beta band (∼15 Hz). These findings suggest that the brain uses an oscillatory mechanism—distinct from previously described beta-bursting inhibitory control—to organize and protect the contents of working memory from inter-item competition.

## Introduction

When multiple items are held in working memory, validly cuing the item that will subsequently be tested has consequences for behavior and for how this information is represented in the brain. Behaviorally, such “retrocuing” results in higher accuracy and faster reaction times, suggesting that the cued item has become “more accessible” (Hautekiet et al., 2025). Neurally, retrocuing improves an item’s decodability (evidence for a “strengthening” of its representation, e.g., Sprague et al., 2016; Yu, Teng, & Postle, 2020), and transforms its representational geometry (Panichello & Buschman, 2021). Intuitively, one might assume that in addition to becoming “stronger,” the cued item also becomes more resistant to interfering effects of other information being processed by the cognitive system, including perceptual distractors. However, empirical evidence for such a “shielding” effect is mixed (e.g., Hautekiet et al., 2025).

Most retrocuing tasks require a single response, thereby rendering the uncued item an “irrelevant memory item” (IMI) that can be “dropped” from working memory once the cue has been processed. For example, in a recurrent neural network (RNN) simulation of the retrocuing task from Panichello & Buschman (2021), the IMI’s manifold contracts, fading from working memory (Piwek et al., 2023). The focus here, in contrast, is a circumstance in which the not-cued item cannot be dropped from working memory because it might be needed later in the trial. One familiar example of such a circumstance is the 2-back task: While item *n* is on the screen, it serves as a recognition probe for item *n* – 2; it next transitions to “unprioritized memory item” (UMI) while *n* + 1 is being compared to *n* – 1; next transitions to “prioritized memory item” (PMI) against which *n* + 2 is compared and finally transitions to IMI. In electroencephalography (EEG) data, the transition from probe to UMI is accompanied by a reversal of the item’s reconstruction by a multivariate inverted encoding model (IEM; Wan et al., 2022). In RNN simulation, the UMI and PMI are represented on ring manifolds that are reversed relative to one another, an item’s geometry undergoing scalar flips during the transition to UMI and then to PMI, before transitioning to IMI and collapsing to a scalar value undifferentiable from 0 (Wan et al., 2022).

The research presented here features a double serial retrocuing (DSR) task in which the presentation of two sample items is followed by a retrocue designating the item that will be tested at the first recall, then a second retrocue, then a second test of recall. As is the case for 2-back (Wan et al., 2022), in DSR the UMI can undergo a representational transformation (Yu, Teng, & Postle, 2020). Furthermore, the transition to UMI has been linked to oscillatory dynamics in the beta band of the EEG: The effects of transcranial magnetic stimulation (TMS) on UMI-linked behavior relates to its effects on beta-band dynamics (Fulvio et al., 2024); and beta-band dynamics make selective contributions to decoding the identity of the UMI (Rose et al., 2016; Fulvio & Postle, 2026). Recent accounts of beta-band dynamics hold that, in a manner analogous to the stopping of skeletomotor actions (Diesburg et al., 2021; Hervault & Wessel, 2025), bursts in the beta frequency range provide a mechanism for imposing top-down control on the processing of information in working memory (Lundqvist et al., 2024). Thus, this report also assesses whether the beta-band dynamics associated with prioritization in the DSR task (Rose et al., 2016; Fulvio et al., 2024; Fulvio & Postle, 2026) may be an example of bursty top-down control (Lundqvist et al., 2024; Wessel & Anderson, 2024).

## Materials & methods

### Experiment 1

#### Participants

Eighteen neurologically healthy members of the University of Wisconsin–Madison community participated in the behavioral experiment. Incomplete data sets from two participants were excluded yielding a final sample of 16 data sets used in the analyses (from 10 females, 2 unknown, 18-32 years, *M* = 22.8 years). The sample size was chosen to match the sample size of Bae & Luck (2017), which reported the inter-item bias effects we aimed to replicate and understand here. All participants had normal or corrected-to-normal vision, and all reported having normal color vision. The research was approved by the University of Wisconsin—Madison Health Sciences Institutional Review Board. All participants gave written informed consent at the start of the session and received monetary compensation in exchange for participation.

#### Experimental procedure

##### Double serial-retrocue task

The double serial-retrocue (DSR; **Fig. 1A**) task began with the 200 ms presentation of an oriented grating at the center of the display (“*Sample 1*”), followed by a 500 ms inter-stimulus fixation interval (“*ISI*”), followed by the 200 ms presentation of a second oriented grating at the center of the display (“*Sample 2*”), followed by 500 ms of fixation (“*Delay 1.1*”), followed by a 500 ms presentation of *Cue 1* (the digit ‘1’ or ‘2’) whose value indicated with 100% validity which of the two samples would be tested. This cue conferred on the cued item the status of the prioritized memory item (PMI) and, by default, the uncued item took on the status of unprioritized memory item (UMI). *Cue 1* was followed by a fixation period (“*Delay 1.2*”) of variable length, densely- and uniformly-sampled across a range from 500-1200 ms in steps of 16.67 ms, implemented in the code as number of screen refreshes. *Delay 1.2* was followed by *Recall 1* in which a circle with a line corresponding to its diameter (and described to participants as a “dial”) was presented. Using the mouse, participants rotated the line to match the orientation of the cued item and clicked the mouse button to lock in their response within a 3-sec window beginning with probe onset. Feedback was displayed for the 1 second interval following the 3 second recall interval, the fixation cross turning green for responses less than 20 deg from the sample orientation, yellow for responses between 20 and 28 deg from the sample orientation, and red for responses greater than 28 deg from the sample orientation. If no response was recorded, the words “No response was recorded” appeared during the feedback interval. The trial then continued with a 500 ms presentation of *Cue 2* (the digit ‘1’ or ‘2’) whose value indicated with 100% validity which of the two samples would be tested during *Recall 2*. The item designated by *Cue 2* took on the status PMI, and the uncued item the status of irrelevant memory item (IMI; because it could no longer be cued on that trial). *Recall 2* and feedback periods were identical to *Recall 1* and its feedback period. After feedback to *Recall 2*, text appeared at the center of the display telling the participant to press the spacebar key when ready to continue with the next trial. This triggered the inter-trial interval, which varied between 2-3 s (“*ITI*”). Upon completion of the block, text appeared on the display informing the participant of their average recall error in degrees on that block. Over the course of each block, *Cue 2* indicated the same item as had *Cue 1* on 50% of trials (“*stay*” trials; the remaining 50% were “*switch*” trials).

##### Neutral-cue task

The neutral-cue (NEU; **Fig. 1B**) task was identical to the DSR task with two exceptions: (i) trials featured only one *Cue* and one *Recall* and *Feedback* period; and (ii) the *Cue* was uninformative, appearing as ‘0’ on each trial. Throughout the text, we will refer to the items in the NEU task not as PMI and IMI, but rather as “tested” and “untested.”

**Fig 1.**
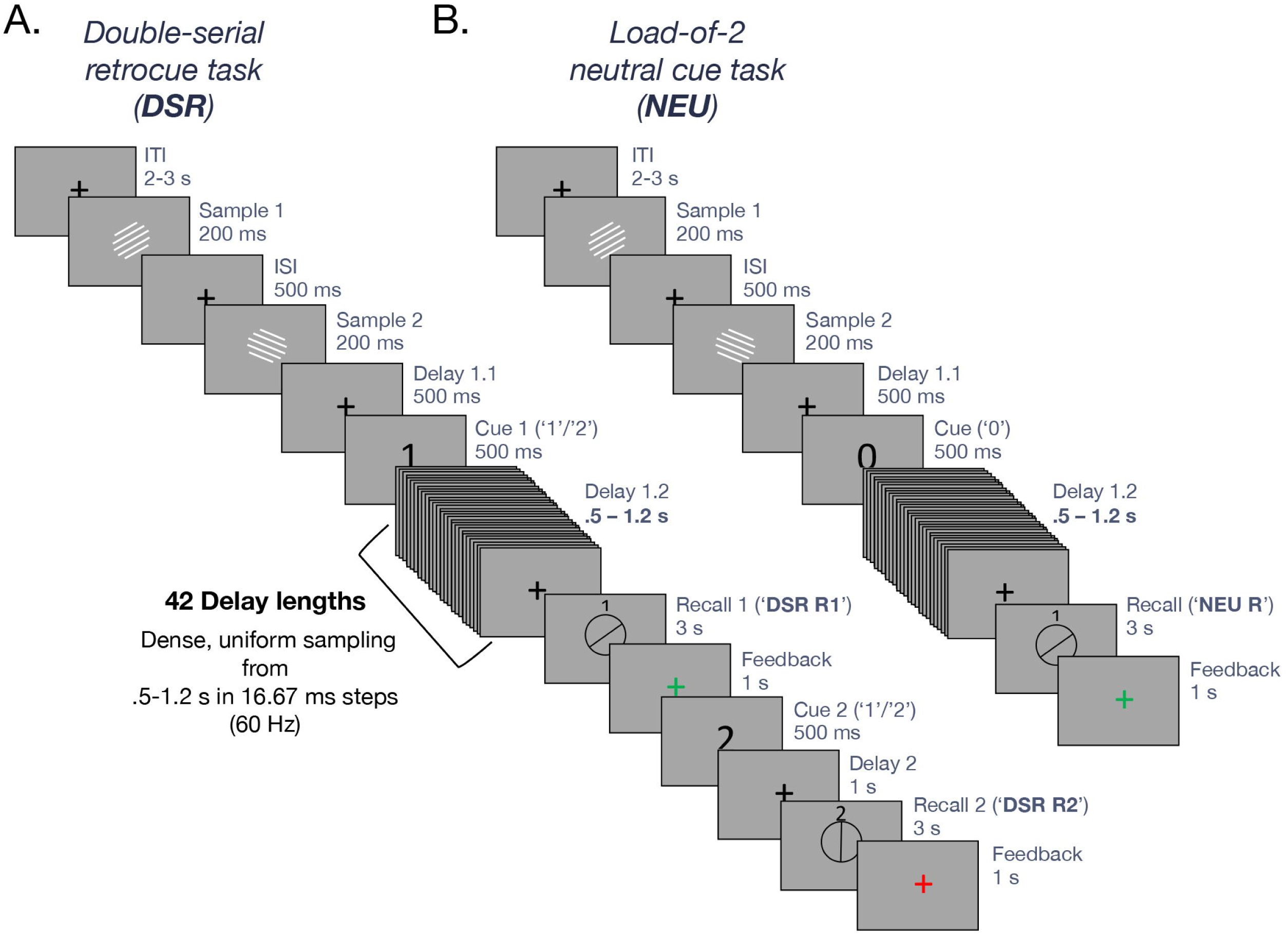
Experiment 1 behavioral task schematics. **A.** Schematic of the double-serial retrocue (DSR) task. Participants memorized the identity of two centrally presented oriented grating samples shown at successive time points with an intervening delay (‘*ISI’*). Partway through the delay (i.e., at the end of *Delay 1.1*) an ordinal retrocue indicated which of the two items would be tested, after the ensuing *Delay 1.2* of variable length, sampled uniformly from .5-1.2 s in 16.67 ms increments. At test (*Recall 1*), participants rotated the recall dial with the mouse to the orientation of the tested sample item in memory. After feedback, a second ordinal retrocue (*Cue 2*) indicating which of the two samples would be tested during *Recall* 2 followed by a final delay. **B.** Schematic of the load-of-2 neutral cue (NEU) task, which followed the same task structure as the DSR task through the first feedback interval, after which the trial ended. Importantly, an uninformative ‘0’ was presented during the cue interval, preventing participants from anticipating which item would be tested at recall.

#### Experimental stimuli and design

Participants were seated in a dimly lit room at a viewing distance of 50 cm from the monitor. Sample stimuli were centrally presented black and white gratings appearing on a mid-gray background with radii of 6°, spatial frequencies equal to 1 cycle/°, 50% contrast, and random phase. The orientation of one of the sample items was selected from a set of six base orientations (10, 40, 70, 100, 130, 160°), with a random jitter of −3° to 3°. The orientation of the other sample item was selected to be offset from the jittered base orientation from a set of inter-item angular differences (0, +/-22.5, +/-45, +/- 67.5°). The presentation order of the base orientation and offset orientation was randomized and counter-balanced across trials. The centrally presented cues were black digits, and the recall stimuli were black response dials (unfilled black circles with a black line corresponding to the diameter of the circle) with the same radius and random starting orientation. A fixation cross was presented during blank task intervals. The cross was gray during the fixation and *ISI* intervals, white during the memory delay periods, and green, yellow, or red during the feedback interval, depending on the response error.

The experiment was completed in a single session, and all participants completed four blocks of the DSR task followed by four blocks of the NEU task. (We reasoned that participants were less likely to ignore DSR cues if they had not previously performed a variant of the task with uninformative cues.) Prior to each set of four blocks, participants completed 8-12 practice trials of the task in the presence of the experimenter. During practice trials, the feedback display differed. After the participant locked in their recall response, the recall stimulus remained on the display and changed color according to the green/yellow/red feedback convention, and the original sample grating reappeared under the recall stimulus to provide visual error feedback to the participant during these trials.

Each block was comprised of 84 trials for a total of 336 trials per task, and participants were encouraged to take breaks between blocks (and between trials if necessary) before continuing. Any trial for which a response was not recorded within the 3-second window was repeated at the end of the block. Because we were primarily interested in behavioral oscillations (and hence the manipulation of the duration of *Delay 1.2*), we established the experimental design to ensure an equal number of trials collected for each of 42 durations of *Delay 1.2*. This resulted in 8 trials collected at each duration, of which there were 2 trials per inter-item angular difference (magnitude) collected per task, per participant. Across those, the base orientations were randomly sampled such that each base orientation was selected for an equal number of trials across all possible base orientation values, but they were not equated across *Delay 1.2* durations.

#### Behavioral analyses

##### Mean absolute error of recall

For both DSR and NEU tasks, we quantified overall accuracy of recall by computing the average magnitude of the angular difference between the participant’s recall response value and the orientation of the tested sample item, as a function of inter-item angular difference and, for the second response of the DSR task, as a function of ‘stay’/‘switch’ identity of *Cue 2*. We also tested for differences in absolute recall error as a function of the target’s presentation order (i.e., as *Sample 1* or *Sample 2*). Paired-sample *t-*tests were used to test for significant differences in mean absolute error of recall across task responses with Bonferroni correction for the number of tests. A linear mixed effects model was used to test for a significant effect of task (2 levels: DSR/NEU, coded as a categorical variable) and inter-item angular difference magnitude (4 levels: 0, 22.5, 45, 67.5 deg). In the model that is reported here, the task x inter-item angular difference interaction term was omitted because this interaction was found to be non-significant and inclusion of the term did not significantly improve the model fit. Subject was included as a random effect, and a separate intercept was fitted for each.

##### Inter-item bias

Following Bae & Luck (2017), we quantified the bias exerted by the untested item on the recall of the tested item as the mean signed recall error, with negative signed errors indicating attraction and positive signed errors indicating repulsion. For each trial, we computed the recall error, and if the response value was in the direction of the untested item, the recall error was given a negative sign, and if the response value was in the direction away from the untested item, the recall error was given a positive sign. As with mean absolute error, we quantified the inter-item bias for both responses during the DSR task and the single response during the NEU task, as well as separately for the second response of the ‘stay’ and ‘switch’ trials of the DSR task, and as a function of inter-sample item angular difference. Paired-sample *t-*tests were used to test for significant differences in inter-item bias across task responses with Bonferroni correction for the number of tests. A linear mixed effects model was used to test for a significant effect of task (2 levels: DSR/NEU, coded as a categorical variable) and inter-item angular difference magnitude (4 levels: 0, 22.5, 45, 67.5 deg) and their interaction with subject included as a random effect and a separate intercept fitted to each.

##### Serial dependence

In addition to within-trial inter-item effects on recall, we also tested for between-trial inter-item effects on recall. For each participant’s data, the response error on each trial was computed as the angular distance between the presented and reported sample orientation. Following previous serial dependence work (Bliss et al., 2017; Fritsche et al., 2017; Samaha et al., 2019; Fulvio et al., 2023), high-error trials (≥25 deg; *M* = 5.4%) were omitted from analysis. To remove any global directional biases of individual participants, we subtracted their mean signed error across trials from each trial. Serial dependence was quantified by sorting each participant’s demeaned errors according to the angular difference between the sample direction on the current and previous trials. Here, we quantified serial bias exerted by the previous trial’s most recent PMI/tested item (i.e., with respect to the PMI designated by *Cue 2* in DSR trials), as well as the bias exerted by the previous trial’s IMI/untested item. In these analyses, the first trial of each block was omitted. For instances in which trials were removed due to large errors, the most immediately preceding non-large error trial served as the previous trial. The sorted errors for each participant were then smoothed by a 20°-wide median filter in 2° increments. (Similar results were obtained with different-sized filters.)

We averaged the smoothed data within each condition and fit the averages of each with a von Mises–based function of the following form:

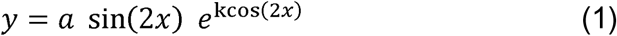

where is the relative orientation of the tested/untested item of the previous trial (in radians), is the amplitude of the curve, and is the concentration parameter determining the width of the tuning. The sine and cosine terms capture the circular periodicity of orientation differences over 180°, allowing the function to model both attractive and repulsive serial biases symmetrically around zero (analogous to a derivative-of-von-Mises (dVM) formulation used in previous studies; e.g., Fritsche et al., 2017). Fitting was achieved by minimizing the sum of squared errors using the *lsqcurvefit* MATLAB routine, with amplitude and concentration parameters free to vary between –10° and 10° and between 0 and 10, respectively.

Statistical significance of the group-level dVM fits was assessed through a bias-corrected and accelerated (BCa) bootstrapping procedure. On each of 100,000 iterations, we sampled participants’ data with replacement and fit a dVM to the average of the bootstrapped sampled data. The value of the amplitude parameter was retained after each iteration, resulting in a distribution of the amplitude parameter of our sample. We also computed the jackknife distribution of amplitude parameters by computing all of the leave-one-sample-out dVM fits. The confidence interval and *p-*value of the amplitudes of each group fit when compared to zero (i.e., no serial dependence) were computed using the BCa_bootstrap MATLAB function (van Snellenberg, 2018), which utilized both the bootstrapped and jackknife amplitude distributions.

Between-condition comparisons of dVM fit amplitudes were carried out using permutation testing. On each of 100,000 iterations, we shuffled the group labels (e.g., ‘DSR’ / ‘NEU’; ‘PMI’ / ‘IMI’) and then assigned each dataset with one of the shuffled labels. We then grouped the data by the shuffled labels and fit a dVM to the average of each shuffled group. The value of the amplitude parameter was retained after each iteration, resulting in a permutation distribution of the amplitude parameter. The *p*-value of the original difference in amplitudes between the two groups was derived from the permutation distribution as the proportion of the permutations greater than the original difference.

##### Linear mixed effects modeling of inter-item effects

After the data were collected, the separate analyses of inter-item influences on recall, as described in the previous section, revealed interference from both current-trial and previous-trial memory items (see **Supplementary Figure 1** for more detail). Because of this, we fit the data from each task to a linear mixed effects model to simultaneously model the inter-item biases of current and previous trial sample items on current trial recall. The dependent variable was the signed recall error on the current trial, modeled as a function of (i.) the orientations of the previous trial’s tested item, (ii.) the previous trial’s untested item, and (iii.) the current trial’s untested item. Specifically, the predictors were derived from the difference between each item’s orientation (in radians) relative to the current trial’s target orientation, computed using the *circ_dist.m* function to ensure proper angular wrapping, and transformed using the sine of twice the angle, sin (2*θ*), to capture orientation-dependent modulations in a directionally signed space. An additional categorical predictor (‘NonTargetPresentation’) coded whether the current trial’s untested item was presented before (1) or after (0) the target item, to assess whether temporal order of presentation modulated inter-item bias. The model took the following form:

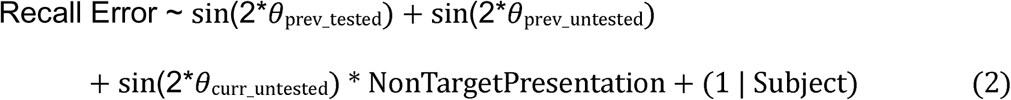

Subject was included as a random effect, with random intercepts to account for inter-individual variability. This model was selected based on nested model comparisons in fitting the NEU data and then used to separately model both the NEU and DSR task *Recall 1* data.

A third model was fit to directly compare the within-trial biases of the two tasks with the form:

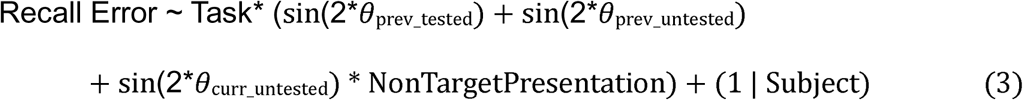

where Task was included as a categorical variable, with the DSR task dummy-coded to be the reference task. This model included the three-way interaction between Task, the current trial’s untested item, and the presentation order of the current trial’s untested item to test whether the effect of prioritization on inter-item bias depended on temporal order of presentation.

After model fitting, we extracted the fixed effects estimates and their associated 95% confidence intervals. Standard errors were computed from the confidence interval widths, and *t*-statistics were obtained as the ratio of each fixed-effect estimate to its standard error. Corresponding two-tailed *p*-values were then derived from the *t* distribution using the model’s residual degrees of freedom. Importantly, positive coefficients for the sine-transformed predictors would indicate attractive biases toward them, whereas negative coefficients would indicate repulsive biases away from them.

To directly test whether the magnitude of inter-item angular difference modulated bias, we conducted a supplementary analysis using a signed bias measure as the dependent variable. This measure was computed by multiplying the signed recall error by the sign of the angular difference between the current trial’s untested and target items, such that positive values indicate attraction toward the untested item and negative values indicate repulsion away from it, maintaining consistency with the primary LME analyses above. The 25% of trials with 0° angular difference were excluded from this analysis because signed bias is undefined when the two items are identical.

The signed bias was modeled as a function of (i.) the angular difference between the current trial’s items (22.5°, 45°, or 67.5°; treated as a categorical variable), (ii.) the presentation order of the current trial’s untested item, and (iii.) their interaction, while controlling for previous trial influences using the sine-transformed orientation differences of the previous trial’s tested and untested items. The model took the following form:

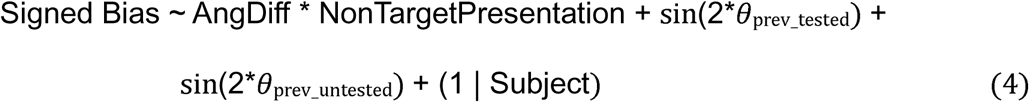

Subject was included as a random effect with random intercepts to account for inter-individual variability. This model was fit separately to the NEU and DSR task data. A third model was fit to directly compare the effects of angular difference across tasks with the form:

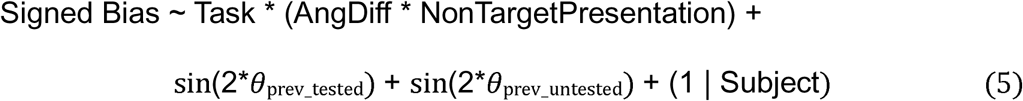

where Task was included as a categorical variable, with the DSR task dummy-coded as the reference level. This model tested whether the effect of angular difference on inter-item bias, and its interaction with presentation order, differed between tasks.

##### Behavioral oscillations

Behavioral oscillations in recall were assessed by analyzing mean absolute error (MAE) as a function of *Delay 1.2* duration (0.5–1.2 s, 42 discrete intervals). For each participant and duration, MAE values were averaged across trials and smoothed with a two-point moving average. The resulting delay-dependent MAE series was linearly detrended to remove slow drifts. To detect rhythmic fluctuations in performance, a Fast Fourier Transform (FFT) was applied to each participant’s detrended MAE time series. Prior to transformation, data were multiplied by a Hann window to reduce spectral leakage and zero-padding to the nearest power of two was applied for computational efficiency and to interpolate the frequency spectrum. The FFT yielded complex coefficients representing amplitude and phase values across 33 frequency bins spanning 0–30 Hz. Group-level spectra were obtained by averaging participants’ complex Fourier coefficients (real and imaginary components), preserving the relative phase information across participants. This approach ensures that consistent phase alignment across participants contributes to the group-level amplitude, rather than being lost through power-only averaging. To assess statistical significance, *Delay 1.2* trial labels were randomly shuffled for each participant, after which delay-dependent MAE was recomputed and the FFT was repeated. This procedure was performed 10,000 times to generate participant-wise null spectra and corresponding group-level amplitude distributions. Observed spectral peaks exceeding the 95th percentile of the null distribution—corrected for multiple comparisons (Bonferroni correction was applied across identified peaks)—were considered significant and would indicate rhythmic modulation of behavioral performance at the frequencies of the significant peaks.

Evidence for participant-level oscillatory structure in the behavioral data was also assessed with a different analysis--an auto-regressive surrogate analysis--in order to provide a second analysis independent of the time-shuffling permutation testing described above. For this, we used the AR-surrogate method of Harris and Beale (2025; see also Fabio & Kayser, 2025; Xu et al., 2026), implemented using the authors’ publicly available MATLAB code (https://github.com/AnthMHarris/BehavOsc-ARSurrogate-PLevel; CC BY 4.0). For each participant, an AR(1) model was fit to their smoothed mean-error time-course via maximum likelihood (Kalman filter;

MATLAB’s arima/estimate), and 5,000 surrogate time-courses were generated from the fitted model. Amplitude spectra were computed via FFT, zero-padded to 64 samples to increase frequency resolution. Consistent with the authors’ implementation, no linear detrending was applied beyond the mean-subtraction step built into their spectral estimation function. A per-participant “deviation” spectrum was calculated as the real amplitude minus the mean surrogate amplitude at each frequency. Group-level significance was assessed with one-tailed one-sample *t*-tests (deviation > 0) at each frequency, which were Holm-Bonferroni corrected across the tested frequency range (θ = .05). Harris and Beale (2025) recommend restricting the analysis to 3–20 Hz based on simulations showing elevated false-positive rates below 3 Hz and above 23 Hz, however, we extended the upper bound to 23 Hz, which was the limit of their reported false-positive control, to encompass the full range of candidate peaks identified in the original permutation analysis.

For frequencies showing significant oscillatory peaks, between-participant phase alignment was quantified using the Phase-Locking Value (PLV). To compute this, each participant’s phase angle at their peak frequency was extracted from the complex FFT output, and PLV was computed as the absolute value of the mean complex phase vector:

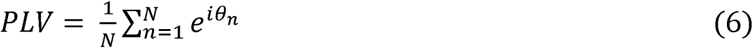

where *N* is the number of participants, θ*_n_* is the phase angle for participant *n*, and *i* =√-1. *PLV* values range from 0 (uniform phase distribution) to 1 (perfect phase alignment). Statistical significance of phase alignment was assessed using both a parametric Rayleigh test for non-uniformity and a non-parametric permutation test. For the latter, *PLV* values were computed for each of the 10,000 permutation-based null datasets, and the observed *PLV* was compared to the resulting null distribution (α=0.05).

All analyses were conducted separately for the DSR and NEU tasks, allowing for direct comparison of peak oscillation frequencies between tasks. Identical procedures were repeated to assess oscillations in within-trial inter-item bias on recall.

### Experiment 2

#### Participants

A separate sample of 16 neurologically healthy members of the University of Wisconsin–Madison community (10 females, 21-28 years, *M* = 23.7 years) participated in the EEG experiment, and all data were used in the final analyses. The sample size was chosen to match the sample size of Experiment 1. All participants had normal or corrected-to-normal vision, and all reported having normal color vision. The research was approved by the University of Wisconsin—Madison Health Sciences Institutional Review Board. All participants gave written informed consent at the start of the session and received monetary compensation in exchange for participation.

#### Experimental stimuli, procedure, and design

The experimental stimuli, procedure, and design were the same as those of Experiment 1, other than the following differences: (i.) *Delay 1.2* of both tasks had a fixed length of 4 s. (ii.) Participants completed four blocks of the DSR task followed by two blocks of the NEU task, with each block consisting of 60 trials. (iii.) A full-factorial design was ensured whereby each inter-item angular difference magnitude was selected for 30 trials, with each base orientation being selected for 5 of those 30 trials yielding 120 total trials of the NEU task. For the DSR task, we doubled the trials to accommodate the ‘stay’/’switch’ aspect of the task, yielding 240 total DSR task trials.

#### EEG recording and preprocessing

EEG was recorded from 63 electrodes with a BrainVision Recorder using active Ag/AgCl electrodes feeding into an ActiCHamp amplifier (BrainProdcuts GmbH). The electrodes were positioned according to the extended International 10–20 system. Impedances were kept below 25 kΩ, and the data were sampled at 1000 Hz. An electrode placed on the forehead served as ground, and all channels were referenced online to channel Fpz and re-referenced offline to the average of all electrodes.

Data were processed offline using EEGLAB (Delorme & Makeig, 2004) with the FieldTrip toolbox (Oostenveld et al., 2011), and custom MATLAB scripts. The data were downsampled to 500 Hz and were bandpass filtered between 1 and 100 Hz with a notch filter centered at 60 Hz. Electrodes exhibiting excessive noise were removed and the remaining electrodes were re-referenced to their average. The continuous data were then epoched to 500 ms prior to *Sample 1* through the feedback in the NEU task, and through the second feedback in the DSR task. Automatic artifact epoch detection and rejection were then performed using the EEGLAB *pop_autorej.m* procedure. The data then underwent dimensionality reduction to 32 principal components, upon which an independent component analysis was applied to identify and remove components containing eye movement artifacts and muscle artifacts using ICLabel (Pion-Tonachini et al., 2019). Finally, previously removed electrodes were interpolated back in using the spherical spline method.

#### Behavioral analyses

We carried out analyses of the mean absolute error of recall and of inter-item bias according to the procedures described in the corresponding sections of the Methods of Experiment 1. Mirroring the reported results of Experiment 1, mean absolute error and grand linear mixed effects modeling of inter-item biases are reported below; see **Supplementary Figure 3** for separate analyses of inter-item bias due to current and previous trial memory items.

#### EEG spectral analysis

##### Broadband oscillatory peak extraction

For both tasks, we isolated oscillatory peaks from *Delay 1.1* and *Delay 1.2* (i.e., the epoch preceding the cue and following the cue), excluding the first 100 ms of both to minimize visual stimulus-related effects. Because *Delay 1.1* was 500 ms in length, this yielded a window of 400 ms from which to extract oscillatory peaks. To match this, analyses of *Delay 1.2* were restricted to a 400 msec-long window that began 100 msec after the offset of the cue. We computed the power spectral density for each epoch using multitaper Fast Fourier Transform (mtmfft) with Hanning tapering across a frequency range of 3-40 Hz. Spectra were calculated for individual trials and then averaged across trials for each participant.

To isolate oscillatory peaks from the background aperiodic (1/f) component, we applied the Specparam algorithm (formerly FOOOF; Donoghue et al., 2020) with the following parameters: 3-40 Hz frequency range; 5-12 Hz peak width limits; 4 peaks per channel maximum; 0.15 minimum peak height; 2.0 peak threshold; fixed aperiodic mode. For each channel, the algorithm fitted the aperiodic component and identified oscillatory peaks. For visualization, we plotted histograms of all identified peaks across participants and channels for each task and delay period. We used two-sample Kolmogorov-Smirnov (K-S) Tests to assess whether the distributions of peak frequencies differed between epochs. Because the K-S test is sensitive to any differences in distribution shape, location, or spread, and detects the largest local discrepancy between distributions, significant results would indicate that there were global distributional changes in oscillatory peak frequencies between epochs. To test for differences in central tendency (median) between the distribution of peaks from the two epochs (i.e., *Delay 1.1* vs. *Delay 1.2*), we used Mann-Whitney U tests, for which significant results would indicate that there were systematic shifts in the peak median frequency between epochs. Finally, we calculated the Wasserstein distance (Earth Mover’s Distance) between the distributions of peaks from the two epochs. This measure quantifies the minimum cost of transforming one distribution into the other, computed across the *entire* frequency distribution, accounting for both the magnitude of differences and the distance over which spectral energy must be displaced (Rubner et al., 2000; Lipp & Vermeesch, 2023). The Wasserstein distance was calculated for the pooled data (i.e., all peaks extracted across participants) between all peak frequencies from *Delay 1.1* and all peak frequencies from *Delay 1.2* using 200 quantile comparison points. We then shuffled the epoch labels and computed the distance 5,000 times and the *p*-value was computed as the proportion of shuffled data sets that showed a distance equal to or greater than the actual observed distance. We also performed a participant-level analysis, computing the distance between epochs for each participant, then shuffling the condition labels within that participant’s data set 2,000 times to create a participant-specific null distribution. To assess whether within-participant distributional shifts were consistent across participants, we randomly sampled one value from each participant’s null distribution and averaged across participants (repeated 2,000 times), and compared the observed group mean distance to this null distribution.

##### Beta band peak extraction (for Hypothesis 1.A.)

We next refined our peak extraction analysis by focusing on the two ranges identified in the MAE behavioral oscillations in Experiment 1: a *low-beta* range of 14-16 Hz; and a *higher-beta* range of 19-21 Hz. The spectral analysis pipeline used for broadband peak extraction was applied to the same *Delay 1.1* and *Delay 1.2* epoch data from the DSR and NEU tasks, extracting peaks only in the specified *low-beta* and *higher-beta* ranges. For each channel and participant, we extracted all peaks in each band and averaged the band counts separately across channels. We then used a three-way repeated measures ANOVA with factors: Task (DSR vs. NEU), Band (*low-beta*: 14-16 Hz vs. *higher-beta*: 19-21 Hz), and Epoch (*Delay 1.1* vs. *Delay 1.2*) to quantify differences in beta peak density (in units of peaks per channel). Follow-up analyses included task-specific paired *t*-tests comparing peak densities across epochs within each band and a focused test of the Band × Epoch crossover pattern identified in the DSR condition. For visualization, we plotted the group-average beta peak density topographically across conditions. These topographic maps are descriptive and were not subject to statistical testing due to sparse peak distributions.

##### Aperiodic component analysis

In addition to analyzing oscillatory peaks, we examined the aperiodic component of the power spectra, which reflects broadband neural activity independent of narrowband oscillations. The Specparam algorithm models the aperiodic component as a 1/f-like function characterized by two parameters: the offset (reflecting broadband power) and the exponent (reflecting the slope of the power spectral decay, which has been proposed as a proxy for the balance between neural excitation and inhibition (“E:I balance”; Gao et al., 2017; Waschke et al., 2021). For each participant, we extracted the aperiodic exponent and offset for each channel, then averaged across channels to obtain a single value per participant for each epoch and task. To test whether aperiodic parameters changed across delay periods, and whether these changes differed by task, we conducted 2 (Task: DSR, NEU) × 2 (Epoch: *Delay 1.1*, *Delay 1.2*) repeated measures ANOVAs separately for offset and exponent. Prior to analysis, we assessed normality of the difference scores using the Lilliefors and Anderson-Darling tests. Because the exponent difference scores violated normality assumptions in the DSR task (Lilliefors *p* = .005, kurtosis = 5.35), we supplemented the parametric ANOVAs with non-parametric Wilcoxon signed-rank tests on the epoch effects collapsed across tasks.

##### Sub-band burst rate analysis (for Hypothesis 1.B.)

To examine frequency-specific temporal dynamics of beta bursts, we separately quantified burst rates in the *low-beta* (14-16 Hz) and *higher-beta* (19-21 Hz) frequency bands. Bursts were detected using the Matlab implementation of the SpectralEvents toolbox (https://github.com/jonescompneurolab/SpectralEvents) with time-frequency analysis conducted via Morlet wavelet decomposition (width = 5) across 13-30 Hz. Following prior studies, findMethod = 1 was used as in Shin et al., 2017, see also Kavanaugh et al. (2026), which is agnostic to event overlap, and an event threshold of 6 x the median power across time and epochs for each frequency bin of the time-frequency representation.

For each participant, bursts with a peak frequency falling within each sub-band of interest were identified, and their rates (events per second) were calculated for the same two temporal epochs used in the oscillatory peak analysis (see ***Broadband oscillatory peak extraction***): the final 400 ms of *Delay 1.1* (pre-cue baseline period) and a 400 ms window beginning 100 ms after cue offset during *Delay 1.2* (post-cue period). Burst rates were averaged across all scalp electrodes to yield a single value per participant, task, epoch, and frequency band.

We first computed two descriptive measures of trial prevalence to characterize burst prevalence. First, we calculated the proportion of trials containing at least one burst within each channel, then averaged these proportions across channels (per-channel prevalence). Second, we calculated the proportion of trials containing at least one burst anywhere across all channels (any-channel prevalence), which provides a more direct estimate of how frequently bursts occurred during the epochs.

Statistical analyses utilized paired *t*-tests comparing low-beta and higher-beta burst *rates* (events/sec) within each task and epoch, paired *t*-tests comparing *Delay 1.1* and *Delay 1.*2 burst rates within each frequency band and task, and separate 2 × 2 repeated-measures ANOVAs (Task [DSR vs NEU] × Epoch [*Delay 1.1* vs *Delay 1.2*]) for each frequency band to assess main effects and interactions.

##### Oscillatory power

To complement the spectral parameterization and burst rate analyses, we examined oscillatory power in the *low-beta* (14-16 Hz) and *higher-beta* (19-21 Hz) frequency bands. Time-frequency decomposition was performed using Morlet wavelets (width = 5 cycles) as implemented in FieldTrip. Power was computed over the timepoints spanning *Delay 1.1* through *Delay 1.2* (a total of 5000 ms sampled at 250 Hz for 1250 timepoints) in 50 ms steps. We decomposed power into induced and evoked components. Total power was computed by first applying wavelet decomposition to each trial and then averaging across trials. Evoked power was computed by first averaging the raw signal across trials to obtain the event-related potential and then applying wavelet decomposition to the averaged signal. Induced power was calculated as total power minus evoked power, isolating oscillatory activity that was not phase-locked to stimulus onset.

Initially, the power values were not baseline-corrected because the period preceding *Delay 1.1* contained stimulus presentation (*Sample 2*) and thus did not provide an appropriate baseline. For statistical analysis, power was averaged across all channels and across the two time windows used in the other spectral analyses described above: the final 400 ms of *Delay 1.1* and the first 400 ms of *Delay 1.2*, offset by 100 ms to avoid sample-evoked and cue-evoked transients, respectively. Induced and evoked power were analyzed separately for each frequency band using 2 × 2 repeated measures ANOVAs with factors Task (DSR, NEU) and Epoch (pre-cue, post-cue). Follow-up paired *t*-tests were carried out to examine simple effects the two factors with a Bonferroni-corrected alpha-level = .025 (for two comparisons per effect/band).

Because strong task differences emerged in the non-baseline-corrected analysis, we carried out a follow-up comparison of the power in the two tasks using *Delay 1.1* as the baseline allowing us to address whether the power in the two tasks exhibited differential cue-related changes in power, rather than just differing in absolute power levels. For each participant, we computed the percent change in power from pre-cue to post-cue as ((*post* − *pre*) / *pre*) × 100 separately for each frequency band and power type (induced, evoked). Paired *t*-tests were carried out to test for differences in percent power change between DSR and NEU at the Bonferroni-correct alpha-level = .025.

#### Priority-related transformation of signals

##### Temporal generalization representational similarity analysis (RSA)

To assess whether the delay-period representational geometry was preserved across the cue transition in the DSR task, we performed a cross-epoch temporal generalization RSA. For each participant and item (PMI, UMI), representational dissimilarity matrices (RDMs) were computed independently within successive, non-overlapping 28-ms (7-sample) windows spanning *Delay 1.1* and, separately, *Cue 1*. Windows were tiled outward from the delay–cue boundary so that no window’s underlying EEG samples spanned both epochs. Each RDM was constructed by condition-averaging the EEG data (63 channels × window samples) across trials within each condition. Conditions were defined by all 24 combinations of inter-item angular difference (0°, +/-22.5°, +/-45°, +/-67.5°) and base orientation (10°–160° in 30° steps), separately for PMIs and UMIs. If there were no trials for a particular combination of angular difference + base orientation in a priority condition (due to the removal of bad trials during pre-processing), that row and column were omitted from the analysis. This occurred for 1 row/column in the analysis of 1 participant’s data. Pairwise dissimilarity was the computed (1 − Pearson correlation) between all condition pairs. Each *Delay 1.1* window’s RDM (vectorized lower triangle) was then correlated (Spearman) with each *Cue 1* window’s RDM, yielding a rectangular delay-by-cue temporal generalization matrix per participant and item. Because the two axes were drawn from non-overlapping epochs, no cell reflects a trivial self- or near-self-comparison. Group-level matrices were computed by Fisher z-transforming each participant’s correlation matrix, averaging across participants, and back-transforming for visualization. To test whether temporal generalization differed by priority, we computed the paired PMI–UMI difference (Fisher z) at each cell and evaluated it with a cluster-based permutation test (1000 permutations, sign-flip null across participants, cluster-forming threshold *p* < .05 two-tailed, cluster-level correction at *p* < .05 based on maximum cluster mass).

##### Representational self-consistency over time

To establish the interpretability of a time-locked fluctuation in a signal that correlates with an event of interest (in our case, a change specific to the cue), it is important to be able to rule out that this fluctuation may be similar to fluctuations that occur at other points in time (i.e., to “general” fluctuations that are not related to the event of interest). To do this, we first assessed the internal self-consistency of the RSA time series within a portion of the *Delay 1.1* epoch and within a portion of the *Cue 1* epoch and then compared these with the self-consistency of this time series within a window that spanned the onset of *Cue 1*. Our approach adopted time x time pattern-similarity approaches that are used to assess representational stability (e.g., Sommer et al., 2022; Pacheco-Estefan et al., 2024). Sommer et al. (2022) developed the general framework for assessing self-consistency by constructing time × time similarity matrices from EEG data by correlating multivariate neural activity patterns across all pairs of time points and using this to assess the temporal stability and distinctiveness of neural representations. They further note that this framework can be extended to representational dissimilarity matrix (RDM) analyses by summarizing pairwise similarities among stimuli into RDMs and comparing those RDMs with one another to assess “second-order isomorphism.” Thus, rather than comparing time × time similarity matrices from EEG (Sommer et al., 2022) or iEEG (Pacheco-Estefan et al., 2024) data, here we compared time × time similarity matrices from the temporal generalization RSA.

Because evaluating representational self-consistency requires comparing RDMs *within* the same epoch, to carry out this analysis we first generated two temporal generalization matrices, one spanning the full 500 ms of *Delay 1.1* and one spanning the full 500 ms of the *Cue 1* epoch on both axes. As in the cross-epoch analysis described above, representational dissimilarity matrices (RDMs) were computed for each participant and condition in successive, non-overlapping 28-ms windows, tiled outward from the delay–cue boundary so that no window spanned both epochs. All pairwise correlations (Spearman) between RDMs within the same epoch were computed, excluding the trivial diagonal (each window correlated with itself). For each participant, these within-epoch correlations were Fisher z-transformed and averaged separately within *Delay 1.1* and within *Cue 1,* yielding one self-consistency value per participant, per epoch, per item (PMI, UMI). Group-level self-consistency was evaluated with one-sample *t*-tests against zero, separately for PMI and UMI in each epoch, and PMI–UMI differences in self-consistency within each epoch were evaluated with paired *t*-tests. Upon establishing statistically significant levels of self-consistency within these epoch-spanning windows (see Results), we repeated this analysis using 112 ms windows positioned immediately before and immediately after the onset of *Cue 1*. The results from these smaller windows, indicating significant self-consistency during the 112 ms prior to *Cue 1* onset as well as during the first 112 ms of *Cue 1* (Results), allowed us to next assess evidence for a *Cue 1*-related influence on brain activity in two ways: by comparing the levels of self-consistency immediately prior to and immediately following cue onset with that of a 112-ms window spanning the onset of *Cue 1* (next), and by comparing the self-correlation of patterns of EEG signals immediately prior to and immediately following cue onset (section *Self-correlation of raw EEG (for Hypothesis 2)*).

To test directly whether the onset of *Cue 1* triggered a shift in the RSA time series of each stimulus item that exceeded what was observed within *Delay 1.1*, and thus plausibly specific to the designation of priority status within working memory, we constructed a single temporal generalization matrix corresponding to a 112-ms window that spanned the delay–cue boundary (four 28-ms windows per side, tiled as in the analyses above) and compared the resultant estimates of “cross-boundary” self-consistency with the estimates of within-epoch self-consistency (i.e., from the final 112 ms of *Delay 1.1* and from the first 112 ms of *Cue 1*. For each participant and item, within-epoch and cross-boundary correlations were Fisher *z*-transformed and averaged separately, and their difference (within-epoch minus cross-boundary) was evaluated with a one-sample *t*-test. A reliable positive difference would indicate that representational structure is more similar within an epoch than across the cue boundary and thereby provide evidence of a specific effect of the cue on stimulus representation. PMI–UMI differences were evaluated with a paired *t*-test on the within-minus-cross difference scores.

One possible complication with interpreting the results from the cross-boundary analysis of self-consistency is that whereas the maximum temporal lag for comparisons between 28-ms windows within each epoch was 84 ms, for cross-boundary comparisons it was 196 ms (i.e., for the earliest *Delay 1.1* window versus the latest *Cue 1* window). Because representational similarity is generally expected to decline as the temporal distance between two windows increases, independent of any other factors (see Results—Representational self-consistency over time), this discrepancy in the average temporal separation within-epoch versus across-epoch created a confound. To isolate effects specific to the cue-onset boundary, we repeated the analysis restricting cross-boundary comparisons to the same 28–84 ms lag range covered by the within-epoch comparisons, so that any remaining difference between within-epoch and cross-boundary correlations could no longer be attributed to a difference in average lag.

##### Self-correlation of raw EEG (for Hypothesis 2)

To examine how neural signals corresponding to prioritized and unprioritized memory items (i.e., PMIs and UMIs) may transform in response to the retrocue, we applied a correlation analysis that operates directly on the raw multichannel voltage patterns (Zhang & Lewis-Peacock, 2026). For each of the 24 conditions that were also used to derive the RDMs for RSA analyses (6 base orientations x 4 inter-item angular differences), we pooled all trials and time samples within the final 112 ms of *Delay 1.1* (i.e., 28 time samples) into a single representative channel-pattern vector and separately did the same for the first 112 ms of *Cue 1*. For each item (PMI, UMI), the correlation between the *Delay 1.1* vector and the *Cue 1* vector was computed for each of the 24 conditions, and the resulting 24 condition-wise correlations were Fisher z-transformed and averaged to yield a single self-correlation value per participant, per epoch, per item. The resulting *Delay 1.1*-vs-*Cue 1* correlation was validated against a bootstrapped ceiling, which was computed by correlating two independent resamples of *Delay 1.1* data against each other, repeated over 500 bootstrap iterations and averaged, to provide an estimate of the correlation expected from trial-sampling noise alone. The “cue-locked drop” was defined as ceiling (Fisher-z) minus the observed correlation (Fisher-z) and indicated how far the observed *Delay 1.1*-to-*Cue 1* similarity fell below what would be produced by pure noise. Paired *t*-tests (Fisher-z scale) comparing the ceiling vs. real *r*, were carried out separately for the PMI and UMI, testing whether each showed a genuine drop below the noise ceiling, and a paired *t*-test on the two items’ drop scores was carried out to test whether the magnitude of the drop differed by item priority status. These analyses were repeated for the NEU task data, and to test for differences between the two tasks, “drop” scores were first collapsed across item type within task (mean of PMI/UMI for DSR; mean of Tested/Untested for NEU) and then compared using a paired *t*-test.

## Results

### Experiment 1

#### Recall accuracy

Mean absolute error (MAE) of recall did not differ between DSR *Recall 1* and NEU recall (*t*(15) = 0.6046, *p* = .555), whereas the MAE of DSR *Recall 2* was significantly larger than for DSR *Recall 1* and NEU recall (both *p*-values < .01; **Fig. 2A**). No difference was observed between the MAE of DSR *Recall 2* for ‘stay’ versus ‘switch’ trials (*t*(15) = 0.0526, *p* = .959; **Supplementary Figure 1A**). Larger inter-item angular differences were associated with larger errors, with a linear mixed effects model confirming a significant main effect of current trial inter-item angular difference on performance (*F*(1,189) = 140.81, *p* < .001; **Fig. 2B**). When the model included both recall responses of the DSR task, a significant main effect of task was also observed (*F*(1,189) = 9.9764, *p* = .002), driven by the worse performance for *Recall 2* of the DSR task (see also **Supplementary Figure 1B**).

**Fig 2.**
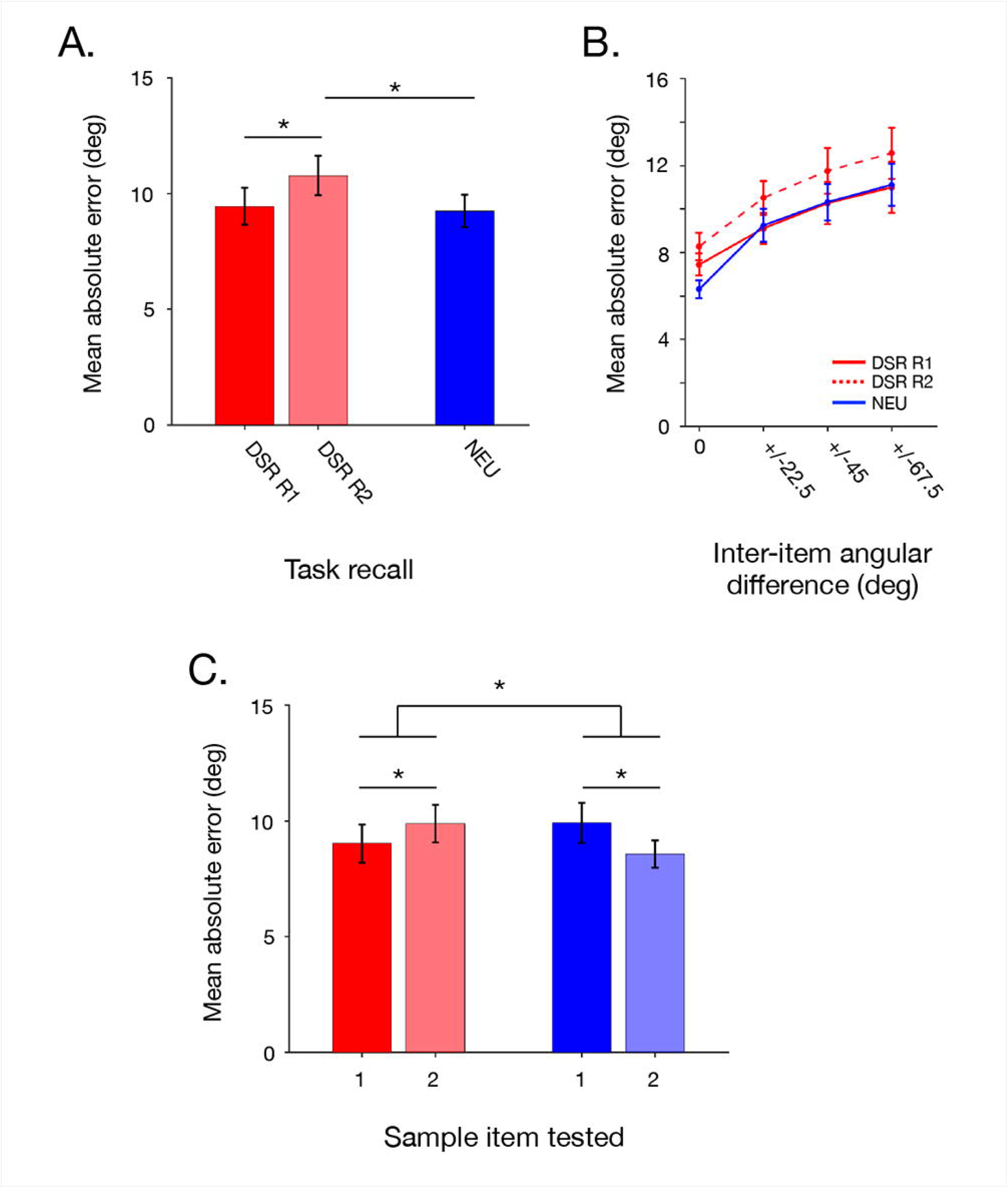
Experiment 1 recall accuracy. **A.** Mean absolute recall error for the two responses of the DSR task and the single response of the NEU task. **B.** Same as **A.** separated by the current trial’s inter-item angular difference. **C.** Mean absolute error for DSR *Recall 1* and NEU recall as a function of which item was tested (*Sample 1* vs. *Sample 2*). Error bars correspond to +/- 1 *SEM.* * symbols correspond to significant differences between conditions at the Bonferroni-corrected alpha level = .0167 for 3 paired-sample *t*-tests (**A.**) and the Bonferroni-corrected alpha level = .025 for two paired-sample *t*-tests and for interaction *p* < .05 (**C.**).

Finally, comparison of MAE as a function of the presentation order of the PMI/tested item revealed different patterns between NEU recall and DSR *Recall 1*: In NEU, MAE was significantly smaller when *Sample 2* was tested (*t*(15) = 2.805, *p* = .0133), whereas in DSR, MAE was significantly smaller when *Sample 1* was tested (*t*(15) = -2.734, *p* = .0154); **Fig. 2C**). This pattern resulted in a significant performance crossover between the two tasks (*t*(15) = -3.849, *p* = .0016).

#### Across- and within-trial bias

Linear mixed effects modeling revealed that response error was systematically influenced by both the previous trial’s tested item (i.e., serial dependence) and the current trial’s untested item (i.e., inter-item bias), with the latter effect dependent on the order of presentation (**Table 1**). In both tasks, the previous trial’s tested item exerted an attractive bias on recall (NEU: β = 1.08°, *p* < .001; DSR: β = 0.84°, *p* = 0.002), an effect that did not differ across task (β = 0.25°, *p* = .517). The previous trial’s untested item had no significant effect on recall. Turning to within-trial effects, recall of *Sample 1* was attracted toward the value of *Sample 2* for NEU (β = 3.55°, *p* < .001) and for DSR (β = 1.66°) but recall of *Sample 2* showed no significant bias in NEU (β = 0.08°, *p* = .818) or DSR (β = 0.50°, *p* = .177). The three-way interaction between task, current-trial untested item, and order-of-presentation was significant (β = –2.31°, *p* = .002), confirming that the effect of prioritization was to reduce the influence of *Sample 2* on the recall of *Sample 1*.

**Table 1.** Linear mixed-effects models were fit separately for the DSR and NEU tasks with identical fixed- and random-effects structures. Only *Recall 1* of the DSR task was used in the full LME analyses. Coefficients are expressed in radians. Coefficients involving sine-transformed predictors and their interactions are also expressed in degrees of response error per unit change in the sine-transformed orientation variable in parentheses. Sine-transformed predictors reflect directionally signed orientation-dependent biases with positive values indicating attractive biasing effects of the predictor and negative values indicating repulsive biasing effects of the predictor. For the current-trial non-target, simple effects are reported separately for trials in which the non-target was presented first versus second, derived from the interaction between presentation order and the non-target orientation term.

| Predictor | DSR Task | NEU Task |
| --- | --- | --- |
| Non-Target Order | $\beta = -0.0029$; 95% CI: [-0.0156, 0.0099]; $p = .66$ | $\beta = 0.0108$; 95% CI: [-0.0016, 0.0232]; $p = .088$ |
| Previous Trial Target ($\sin 2\theta$) | $\beta = 0.0146$ ($0.84^\circ$); 95% CI: [0.0053, 0.0239]; $p = .002$ | $\beta = 0.0189$ ($1.08^\circ$); 95% CI: [0.0098, 0.028]; $p < .001$ |
| Previous Trial Non-Target ($\sin 2\theta$) | $\beta = -.0086$ ($-0.49^\circ$); 95% CI: [-0.0179, 0.0007]; $p = .07$ | $\beta = -.0032$ ($-0.18^\circ$); 95% CI: [-0.0123, 0.0059]; $p = .49$ |
| Current Trial Non-Target ($\sin 2\theta$) | | |
| --non-target presented second | $\beta = 0.0289$ ($1.65^\circ$); 95% CI: [0.0159, 0.0418]; $p < .001$ | $\beta = 0.0620$ ($3.55^\circ$); 95% CI: [0.0496, 0.0744]; $p < .001$ |
| --non-target presented first | $\beta = .0087$ ($0.50^\circ$); 95% CI: [-0.0039, 0.0213]; $p = .18$ | $\beta = .0015$ ($0.08^\circ$); 95% CI: [-0.0109, 0.0138]; $p = .82$ |
| Non-Target Order x Current Trial Non-Target | $\beta = -0.0202$ ($-1.16^\circ$); 95% CI: [-0.0382, -0.0021]; $p = .029$ | $\beta = -0.0606$ ($-3.47^\circ$); 95% CI: [-0.0781, -0.043]; $p < .001$ |

Linear mixed effects modeling further revealed that inter-item angular difference significantly modulated bias (*F*(2,3977) = 7.16, *p* < .001) in the NEU task, and this effect interacted with order of presentation (*F*(2,3977) = 6.35, *p* = .002). Recall of *Sample 1* was significantly attracted toward *Sample 2* at small and medium angular separations (22.5°: 3.68°, *p* < .001; 45°: 3.72°, p < .001), with attraction reduced to a marginally significant level at the large angular separation (67.5°: 1.09°, p = .052; a significant reduction from the bias at 22.5: β = -2.59°, *p* = .001). Recall of *Sample 2*, in contrast, showed no significant bias relative to *Sample 1* at any angular separation (22.5°: -0.42° *p =* .461; 45°: 0.01° *p* = .987; 67.5°: 0.63° *p* = .251). The main effect of presentation order was significant (*F*(1,3977) = 26.72, *p* < .001). In DSR, bias was small and the pattern of bias did not vary significantly across inter-item angular differences (*F*(2,3975) = 0.70, *p* = .499) nor was there an interaction of inter-item angular difference interaction with order of presentation (*F*(2,3975) = 1.65, *p* = .192). Recall of *Sample 1* was attracted to *Sample 2* at medium and large angular separations (45°: 1.76°, *p* = .002; 67.5°: 1.42°, *p* = .013) but not at the small separation (22.5°: 0.81°, *p* = .155). Recall of *Sample 2*, in contrast, showed no significant bias relative to *Sample 1* at small and medium angular separations (22.5°: -0.71°, *p* = .208; 45°: 0.22°, *p* = .698), but attraction at the large angular separation (67.5°: 1.67°, *p* = .003). Importantly, the previous trial’s target, which exerted a significant attractive bias on recall in the primary analysis, did not significantly influence signed bias in either task. This indicates that the previous trial’s influence operated separately in absolute orientation space, pulling responses toward the previous target’s orientation, rather than by influencing interactions between the current trial’s items.

The task comparison model confirmed that NEU showed significantly greater overall bias than DSR (*F*(1,7954) = 12.98, *p* < .001), and that the effect of angular difference was significantly larger in NEU than DSR (interaction: *F*(2,7954) = 4.21, *p* = .015).

#### Behavioral oscillations

Permutation-based FFT analysis revealed significant rhythmic fluctuation of mean absolute error, at different frequencies, on both tasks (**Fig. 3A**). DSR showed significant behavioral oscillatory peaks clustered around 15 Hz (14.06, 15.00, and 15.94 Hz), with the dominant frequency at 15.00 Hz (amplitude = 0.34 degrees). The NEU task exhibited significant behavioral oscillations with the dominant frequency at 20.62 Hz (amplitude = 0.18 degrees; range = 19.69 and 20.62 Hz). We additionally applied the participant-level AR-surrogate method proposed by Harris & Beale (2025) to assess the significance of the peaks in the mean absolute error traces against a null distribution that preserves the aperiodic (1/f-like) structure of each participant’s time-course. Data from the DSR task showed significant oscillatory power at two of the three frequencies that were identified by the time-shuffling procedure: 14.06 Hz (*t*(15) = 3.41, *p* = .040 corrected) and 15.00 Hz (*t*(15) = 3.31, *p* = .048 corrected; for the 15.94 Hz peak, its *p =* .0134 did not survive correction (*p =* .2414 corrected)). For the NEU task, the two frequencies identified by the time-shuffling procedure did not survive correction using the AR approach: 19.69 Hz: *p =* .0281, *p =* .4771 corrected; 20.62 Hz: *p =* .2363, *p =* 1 corrected. These results thus provide confirmatory evidence for a low-beta oscillation specific to prioritization in the DSR; but leave equivocal whether cuing with a neutral cue may enhance behavioral oscillation in a “higher-beta” frequency band.

Participant-level peak frequency analysis confirmed that the peak oscillatory frequencies differed significantly between conditions (*M* = 5.15 Hz; *95% CI*: [3.38, 6.93] Hz; *t*(15) = -6.18, *p* < 0.001). From here forward, to relate them to the classical frequency band of EEG oscillation, we will refer to these two frequencies as “low-beta” and “higher-beta,” respectively.

**Fig 3.**
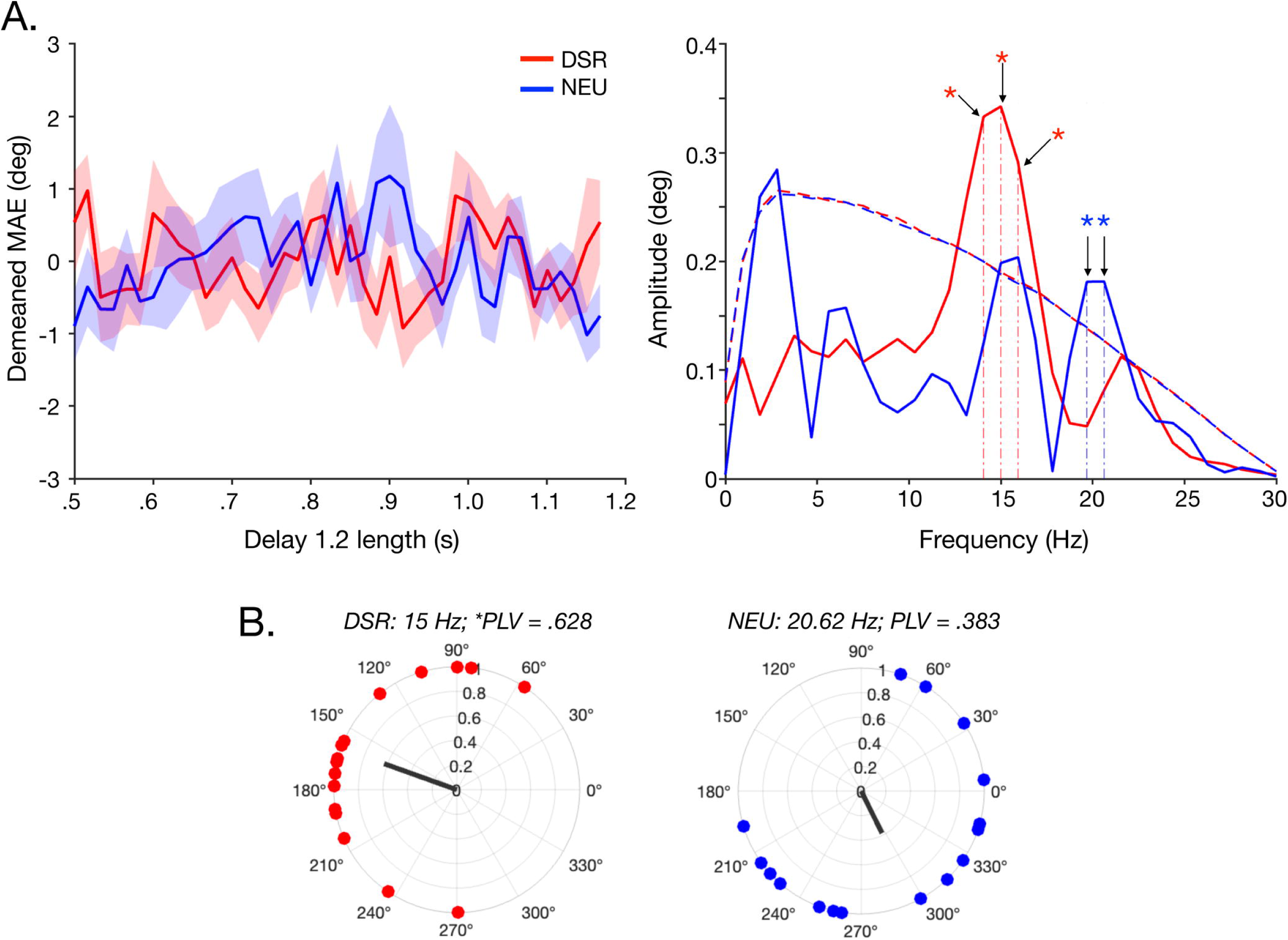
Experiment 1 behavioral oscillations of mean absolute error. **A.** Demeaned mean absolute recall error for DSR R1 (red) and the single response of the NEU task (blue) as a function of Delay 1.2 length. Error bands correspond to +/- 1 *SEM.* **B**: Fast Fourier Transform (FFT) amplitude spectra of mean absolute error across delay periods for the DSR (red) and NEU (blue) tasks. Solid lines represent group-averaged amplitudes (in degrees of error). Dashed lines show statistical significance thresholds (95^th^ percentile) derived from permutation testing. Vertical dot-dashed lines and * symbols indicate peaks surviving Bonferroni-correction for number of peaks exceeding statistical threshold. **C.** Polar plots depicting phase angles at the dominant oscillatory frequency in the DSR task (15.00 Hz; left) and the NEU task (20.62 Hz; right). Individual participants are shown as dots at unit radius. The black line represents the mean resultant vector, with its length corresponding to the Phase Locking Value (PLV) and direction indicating the mean phase angle. * denotes significant PLV.

Between-subject phase alignment was quantified for the dominant frequency of each task. In the DSR task, participants exhibited phase locking of performance fluctuations at 15 Hz (Phase Locking Value [PLV] = 0.628, permutation *p* = 0.0008). In the NEU task, between-subject phase alignment was not significant at 20.62 Hz (PLV = 0.383, permutation p = 0.095), reflecting weaker and less consistent phase coherence across participants. Rayleigh tests for non-uniformity confirmed these findings (DSR: *z* = 6.31, *p* = 0.001; NEU: *z* = 2.34, *p* = 0.095; **Fig. 3B**). To test whether between-subject phase alignment differed between the tasks, we conducted permutation tests comparing PLVs. When comparing each task at its dominant frequency (DSR at 15 Hz vs. NEU at 20.62 Hz), we found that the PLV difference was not significant (Δ*PLV* = 0.245, *p* = 0.33). Thus, while DSR exhibited significant between-subject phase alignment at 15 Hz and NEU did not at 20.62 Hz, the magnitude of phase locking did not differ reliably between tasks.

When we applied the FFT analysis to inter-item bias, no significant oscillatory peaks were observed in either task (all *p* > 0.05), indicating that, unlike MAE, inter-item bias did not fluctuate rhythmically (**Supplementary Figure 2**).

#### Interim discussion—Experiment 1

The results from Experiment 1 provided support for three effects of prioritization on working memory. First, prioritization in the DSR task was associated with a reduction in inter-item interference at recall: The biasing influence of *Sample 2* on the recall of *Sample 1* was markedly reduced. Furthermore, although small biases were present at some angular separations, a critical difference from the NEU task was that for DSR, biases were uniform across separation magnitudes, whereas for NEU, attraction was strong when items were similar and reduced when items were dissimilar. This suggests that prioritization has a general effect of attenuating inter-item interference, reducing sensitivity to the similarity between items. Second, prioritization altered the effect of presentation order on recall accuracy. In NEU, recall was more accurate for the more recently presented item (*Sample 2*), consistent with recency advantages observed in prior multi-item working memory studies (Vergauwe et al., 2016; Pomper & Ansorge, 2021; Chota et al., 2022). Although the bases of this recency effect are not fully understood, one possible explanation is that the most recently presented item may, by default, be most likely to be retained in an encoding-related focus of attention (c.f., Vergauwe et al., 2016). In DSR, in contrast, this pattern was reversed: when *Sample 1* was prioritized, it was recalled more accurately than when *Sample 2* was prioritized. This suggests that prioritization is particularly effective for the earlier-presented item, likely due, in part, to the reduction of the otherwise disproportionate influence exerted by the most recently presented item. Third, prioritization in the DSR task was associated with a lower frequency of behavioral oscillation of mean absolute error (15 Hz; “low-beta”) relative to the NEU task (20.62 Hz; “higher-beta”). Furthermore, only in DSR was the cue associated with phase-locking of the behavioral oscillation across participants (although this effect did not differ significantly between the two tasks). Coupled with the absence of evidence of rhythmic modulation of the strength of inter-item bias, these results suggest that behavioral oscillations of MAE reflect fluctuations in representational strength rather than periodic distortions in the remembered content. This evidence for behavioral oscillations in the beta band is of particular interest in light of previous work demonstrating an important role for beta-band dynamics in the encoding of priority in working memory (Rose et al., 2016; Fulvio et al., 2024; Fulvio & Postle, 2026). For these reasons, Experiment 2 was carried out to test explicitly the hypotheses that oscillatory dynamics in *low-beta* and in *higher-beta* would be associated with the processing of the PMI and the UMI, respectively.

Rhythmicity is a prominent feature of neural systems supporting visual attention (Buschman & Miller, 2009; Fiebelkorn & Kastner, 2019) and visual working memory (Lundqvist et al., 2016; Bastos et al., 2018; Abdalaziz et al., 2023). Experiment 1’s evidence for beta-clocked behavioral oscillations highlight an important question about EEG dynamics observed during working memory: To what extent is this activity reflective of one or more truly periodic processes versus activity that, when assessed at the level of the single trial, is fundamentally bursty. In skeletomotor control, “beta bursts” have a prominent role in stopping the execution of motor actions (e.g., Diesburg et al., 2021; Hervault & Wessel, 2025) and, by analogy, this phenomenon has been proposed as a mechanism for imposing inhibitory control across many domains of high-level cognition, including long-term memory, visual attention, visual working memory, and language processing (Lundqvist et al., 2024; Wessel & Anderson, 2024). Empirically, reanalysis of a dataset originally interpreted as showing sustained, load-dependent activity in the alpha and beta bands during the delay period of a working memory task has revealed that, at the single-trial level, this activity was driven by oscillatory bursts (Kavanaugh et al., 2026). Despite these considerations, however, it is difficult for us to conceive of how a relatively sparse, stochastic, and bursty phenomenon could underlie a behavioral oscillation derived from a time series that was assembled via the after-the-fact sorting of trials into order of duration of the delay period. Thus, an important question for Experiment 2 was to assess whether, at the single-trial level, activity in the *low-beta* and *higher-beta* frequency bands of the EEG would more closely resemble persistent oscillating signals or stochastic bursting.

### Experiment 2

In order to test specific hypotheses relating to the idea that oscillatory dynamics in the *low-beta* band underlie priority-related effects in working memory, we recorded the EEG while participants performed versions of the DSR and NEU tasks that differed from Experiment 1 only in that a single fixed duration was used for *Delay 1.2*. Specifically, we planned to test the following a priori predictions that followed from the results of Experiment 1. First (*Hypothesis 1.A.),* in DSR, the onset of *Cue1* would trigger an increase of activity in *low-beta* but not in *higher-beta*, whereas in NEU, there would be no comparable cue-locked increase in *low-beta.* (Note, however, that for NEU we did not have a strong prediction about whether *Cue1* would be associated with no change in *higher-beta*, or perhaps an increase in *higher-beta*, due to the discrepancy in results between the two analyses of evidence for behavioral oscillation; for each of these tests, changes in activity would be operationalized as the number of oscillatory peaks identified by an analysis that decomposes the EEG into periodic (i.e., “true oscillations”) versus aperiodic components). Relatedly, and assuming interpretable results from *Hypothesis 1.A*, *Hypothesis 1.B* would test the proposition that the task-related dynamics observed in the analyses related to *Hypothesis 1.A*. are better characterized, at the single-trial level, as a consequence of a bursty process (rather than of an underlying periodic signal). Second (*Hypothesis 2*), a larger prioritization-related transformation would be observed for the UMI, rather than for the PMI, because previous studies have observed larger prioritization-related changes in decoding for the UMI (Yu, Teng, & Postle, 2020; Wan et al., 2022). This hypothesis would be operationalized via a self-correlation analysis assessing the degree of cue-related changes in whole-topography EEG patterns. (Additionally, and of tertiary interest to these studies, but addressing previous work from Abdalaziz et al., (2023)), we include in Supplemental Material a test of *Supplemental Hypothesis 3*—the prediction that cue-triggered prioritization interacts with a beta-clocked phase encoding scheme that serves to shield the two memory items from each other.)

#### Behavioral performance

##### Recall accuracy

Overall levels of mean absolute error (MAE) of recall were comparable to what was observed in Experiment 1: MAE did not differ statistically between DSR *Recall 1* and NEU recall (*t*(15) = -2.0024, *p* = .064), nor for DSR *Recall 2* versus NEU recall (*t*(15) = 2.3483, *p* = .033 (Bonferroni-corrected = .0167); **Fig. 4A**). DSR *Recall 1* was superior to DSR *Recall 2* (*t*(15) = 4.7537, *p* < .001), and ‘stay’ and ‘switch’ trials from DSR *Recall 2* did not differ (*t*(15) = -1.5845, *p* = .134; **Supplementary Figure 3**). A linear mixed effects (LME) model confirmed that larger inter-item angular differences were associated with larger absolute errors (main effect: *F*(1,189) = 57.837, *p* < .001; **Fig. 4B**).

**Fig 4.**
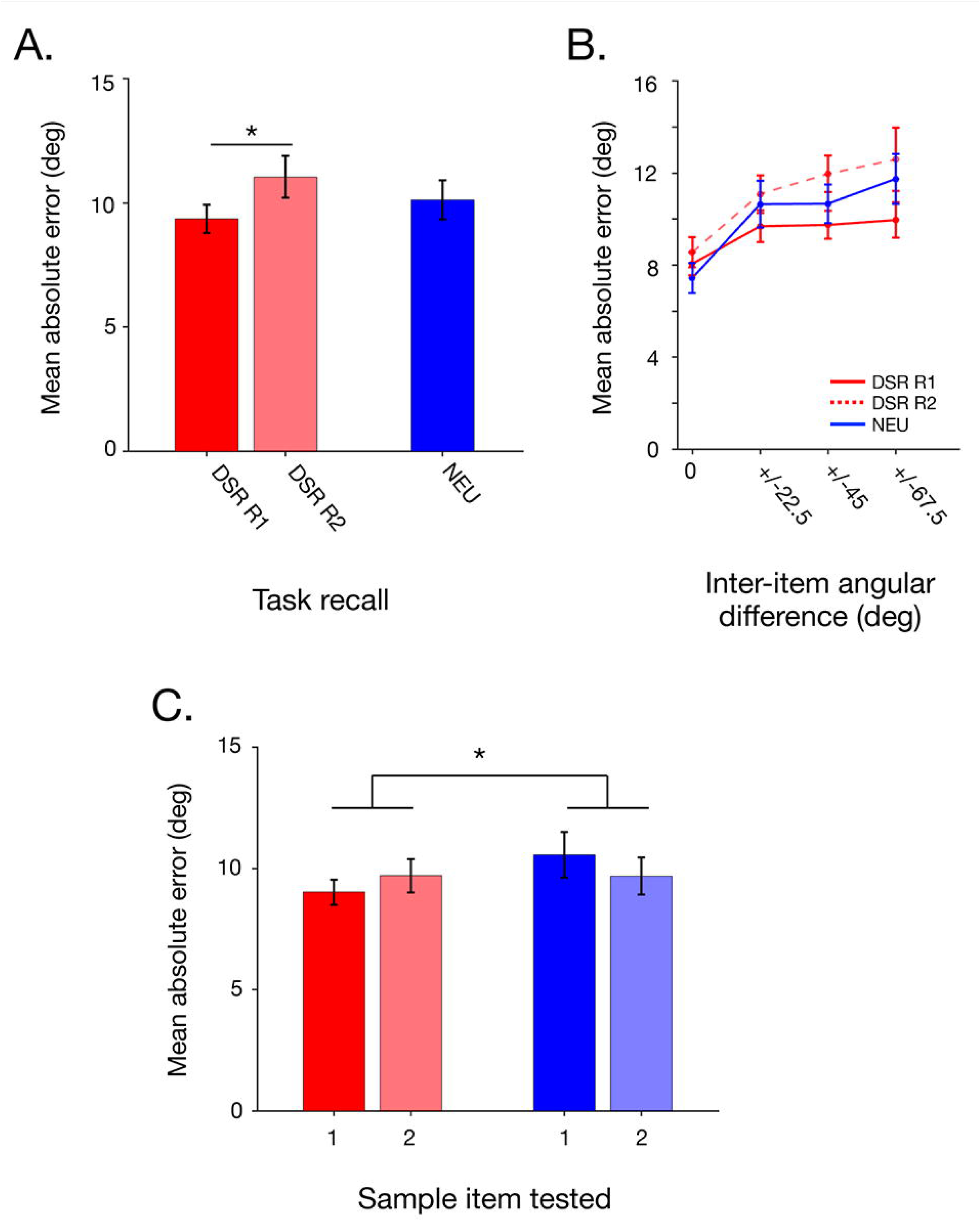
Experiment 2 recall performance. **A.** Mean absolute recall error for the two responses of the DSR task and the single response of the NEU task. **B.** Same as **A.** separated by the current trial’s inter-item angular difference. **C.** Mean absolute error for DSR *Recall 1* and NEU recall as a function of which sample item was tested. Error bars correspond to +/- 1 *SEM.* * symbols correspond to significant differences between conditions at the Bonferroni-corrected alpha level = .0167 for 3 paired-sample *t*-tests (**A.**) and the Bonferroni-corrected alpha level = .025 for two paired-sample *t*-tests and for interaction *p* < .05 (**C.**).

Assessment of MAE as a function of the presentation order indicated that, for NEU, MAE was numerically larger when *Sample 1* was tested (*t*(15) = 1.204, *p* = .247), for DSR, MAE was numerically smaller when was tested (*t*(15) = -1.593, *p* = .132; **Fig. 4C**), and the crossover interaction between the two was significant (*t*(15) = -2.276, *p* = .038).

##### Across- and within-trial bias

Linear mixed effects modeling indicated that the previous trial’s tested item exerted a significant attractive bias on the current trial (NEU: β = 1.60°, *p* < .001; DSR: β = 0.63°, *p* = 0.04; **Table 2**), an effect that was larger in the NEU task (β = 0.97°, *p* = .032). The previous trial’s untested item had no significant effect on recall in either task. Within-trial effects were dependent on the order of presentation. The recall of *Sample 1* was attracted toward the value of *Sample 2* for NEU (β = 4.10°, *p* < .001), but not for DSR (β = -0.39°, *p* = .364), whereas recall of *Sample 2* showed no bias from *Sample 1* in NEU (β = -0.72°, *p* = .131) but a significant repulsive bias in DSR (β = -1.52°, *p* < .001; **Table 2**). The three-way interaction between task, current untested item, and order-of-presentation was significant (β = -3.67°, *p* < .001).

**Table 2.** Linear mixed-effects models were fit separately for the DSR and NEU tasks with identical fixed- and random-effects structures. Coefficients are expressed in radians. Coefficients involving sine-transformed predictors and their interactions are also expressed in degrees of response error per unit change in the sine-transformed orientation variable in parentheses. Sine-transformed predictors reflect directionally signed orientation-dependent biases with positive values indicating attractive biasing effects of the predictor and negative values indicating repulsive biasing effects of the predictor. For the current-trial non-target, simple effects are reported separately for trials in which the non-target was presented first versus second, derived from the interaction between presentation order and the non-target orientation term. Compare to Table 1.

| Predictor | DSR Task | NEU Task |
| --- | --- | --- |
| Non-Target Order | $\beta = -0.005$; 95% CI: [-0.019, 0.0095]; $p = .508$ | $\beta = -0.003$; 95% CI: [-0.019, 0.013]; $p = .749$ |
| Previous Trial Target ($\sin 2\theta$) | $\beta = 0.011$ (0.63°); 95% CI: [0.0052, 0.0217]; $p = .039$ | $\beta = 0.028$ (1.60°); 95% CI: [0.017, 0.040]; $p < .001$ |
| Previous Trial Non-Target ($\sin 2\theta$) | $\beta = -0.010$ (-0.57°); 95% CI: [-0.021, 0.0003]; $p = .058$ | $\beta = -0.004$ (-0.23°); 95% CI: [-0.015, 0.008]; $p = .515$ |
| Current Trial Non-Target ($\sin 2\theta$) | | |
| --non-target presented second | $\beta = -0.007$ (-.39°); 95% CI: [-0.021, 0.008]; $p < .364$ | $\beta = 0.072$ (4.10°); 95% CI: [0.056, 0.088]; $p < .001$ |
| --non-target presented first | $\beta = -0.027$ (-1.52°); 95% CI: [-0.041, -0.012]; $p < .001$ | $\beta = -0.013$ (-0.72°); 95% CI: [-0.028, 0.004]; $p = .13$ |
| Non-Target Order x Current Trial Non-Target | $\beta = -0.0197$ (-1.23°); 95% CI: [-0.04, 0.0006]; $p = .058$ | $\beta = -0.084$ (-4.81°); 95% CI: [-0.107, -0.061]; $p < .001$ |

Linear mixed effects modeling further revealed that in the NEU task, the effect of inter-item angular difference on bias was marginal (*F*(2,2828) = 2.57, *p* = .077), but inter-item angular difference significantly interacted with order of presentation in recall bias (*F*(2,2828) = 3.94, *p* = .019). Recall of *Sample 1* was significantly attracted toward *Sample 2* at all angular separations (22.5°: 4.56°, *p* < .001; 45°: 3.21° *p <* .001; 67.5°: 2.30° *p* = .002) with a significant reduction at the largest angular separation (β = -2.26°, *p* = .025). Recall of *Sample 2,* in contrast, exhibited significant repulsion from *Sample 1* at the small angular separation (22.5°: -1.72° *p* = .021) but no significant bias at medium and large angular separations (45°: 0.01° *p* = .986; 67.5°: -0.20° *p =* .809; main effect of presentation order: *F*(1,2828) = 37.99, *p* < .001).

In the DSR task, there was a significant main effect of presentation order (*F*(1,2827) = 6.74, *p* = .009) and a significant interaction between presentation order and inter-item angular difference (*F*(2,2827) = 4.67, *p* = .009). Recall of *Sample 1* showed no significant bias relative to *Sample 2* at any angular separation (22.5°: -0.01° *p* = .992; 45°: -0.40° *p =* .535; 67.5°: -0.57° *p* = .387). Recall of *Sample 2,* in contrast, was significantly repulsed from *Sample 1* at small and medium angular separations (22.5°: -2.33° *p* < .001; 45°: -2.03° *p* = .001) but no bias at large separations (67.5°: 0.75° *p* = .220; interaction: β = 3.64°, *p* = .004).

The previous trial’s target did not significantly influence signed bias in either task (DSR: *p* = .288; NEU: *p* = .157). The task comparison model confirmed that NEU showed significantly greater overall bias than DSR (*F*(1,5657) = 24.49, *p* < .001), and that the effect of presentation order on bias differed significantly between the tasks (interaction: *F*(1,5657) = 8.22, *p* = .004): in NEU, presenting the untested item first eliminated attraction, whereas in DSR, it induced repulsion. The effect of inter-item angular difference did not differ significantly between tasks (interaction: *F*(2,5657) = 0.69, *p* = .502).

In most regards, these results replicated the pattern from Experiment 1 that prioritization shielded the cued item from the influence of other stimuli. In particular, the attractive bias toward the previous trial and the attractive bias of *Sample 2* on recall of *Sample 1* were markedly smaller in DSR than in NEU. The angular difference analysis further revealed that, as in Experiment 1, the previous trial’s target did not influence signed bias, confirming that between-trial and within-trial sources of recall bias operate independently. However, several other findings diverged from Experiment 1. First, recall of *Sample 2* showed greater repulsion from *Sample 1* in DSR vs. NEU. Second, for DSR, recall of *Sample 2* showed repulsion from *Sample 1* for angular separations of 22.5° and 45°. Third, for NEU, recall of *Sample 1* showed a uniform pattern of attraction across angular separations and a significant repulsion of *Sample 2* from *Sample 1* for angular separations of 22.5°. We will discuss these effects in Interim Discussion— Experiment 2.

#### EEG spectral analysis

##### Oscillatory components

(Test of Hypothesis 1.A.) For the DSR task, visual inspection of the full distribution of extracted peak frequencies pooled across all participants and channels revealed a post-cue increase in the number of peaks in the *low-beta* range (**Fig. 5A**), an effect confirmed by K-S test (*D* = .1367, *p* < .001). A follow-up Mann–Whitney U test found that the difference identified by the K-S test was not due to a shift in central tendency (*z* = 1.13*, p* = .260), indicating that the effect reflected differences in distributional shape. This was confirmed by a permutation test on Wasserstein distance between the two pooled distributions, which was significant (mean distance = 1.22 Hz, permutation *p* < .001). To assess whether these pooled differences were consistent across participants, we computed the Wasserstein distance between each participant’s peak distributions in the two epochs. Participants consistently exhibited significant non-zero within-subject distributional differences (mean distance = 2.59 Hz; permutation *p* < .001).

**Fig. 5.**
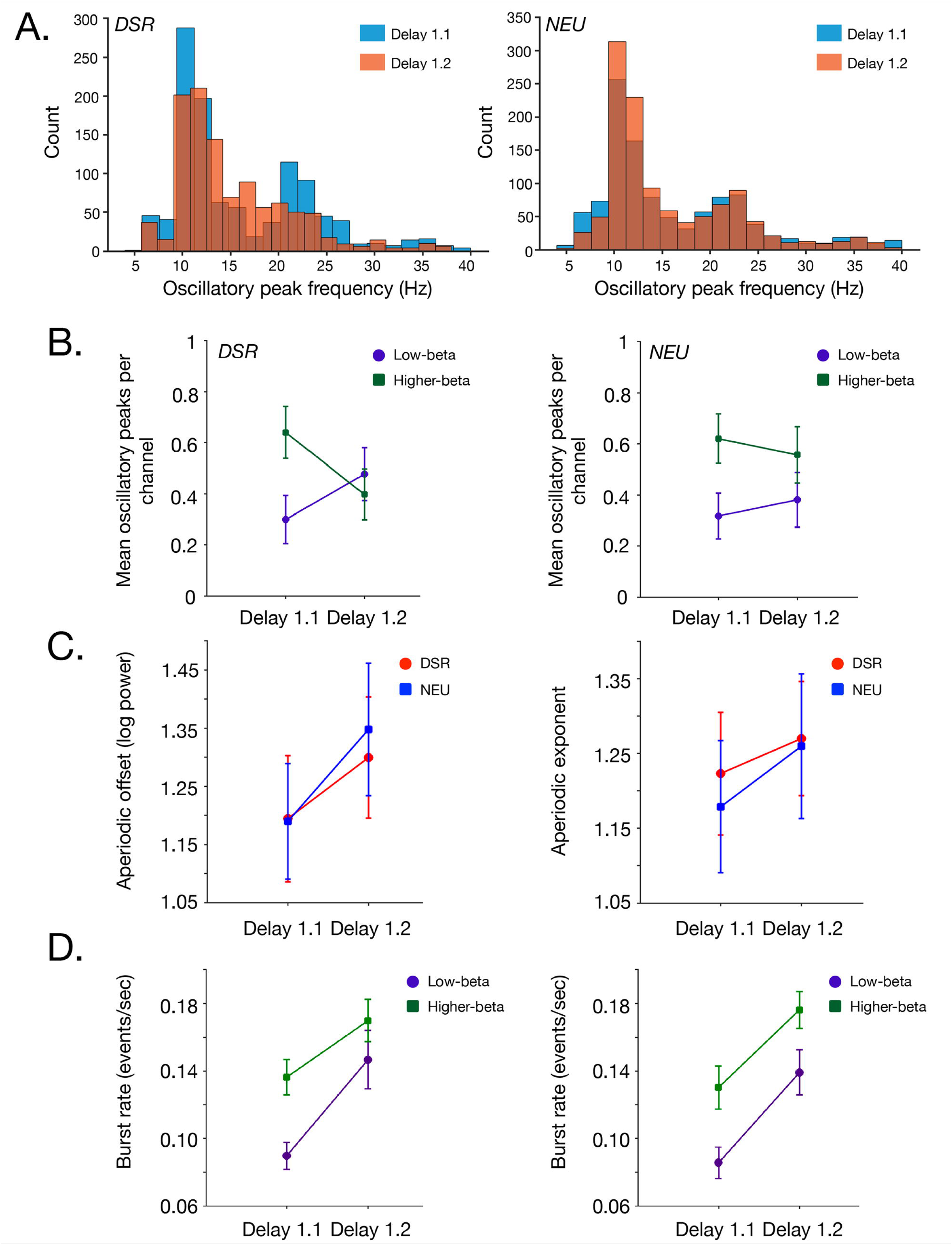
EEG spectral properties. **A.** Distribution of all oscillatory peak frequencies identified across channels and participants from *Delay 1.1* (light blue) and *Delay 1.2* (orange) for the DSR task (left) and for the NEU task (right). **B.** Group-averaged mean number of low beta peaks (indigo circles) and higher-beta peaks (green squares) per channel from *Delay 1.1* and *Delay 1.2* for the DSR task (left) and the NEU task (right). **C.** Left: Mean aperiodic offset for *Delay 1.1* and *Delay 1.2* in the DSR task (red circles) and in the NEU task (blue squares). Right: Mean aperiodic exponent for *Delay 1.1* and *Delay 1.2* in the DSR task (red circles) and in the NEU task (blue squares). **D.** Group-averaged mean low-beta burst rate (indigo circles) and higher-beta burst rate (green squares) per channel from *Delay 1.1* data and *Delay 1.2* data for the DSR task (left) and the NEU task (right). Error bars correspond to +/- 1 *SEM*.

Results for the NEU task differed from DSR: the cue-locked change of the full distributions of peak frequencies was a shift toward a greater number of peaks in the alpha frequency range (**Fig. 5A**), an effect confirmed by K-S test (*D* = .0623, *p* = .024), with no change in median peak frequency (*z* = 0.1675*, p* = .867). At the group level, participants consistently exhibited non-zero Wasserstein distances between epochs (mean distance = 2.31 Hz; permutation *p* < .001), indicating that the distributions of peaks in the NEU task also changed in a consistent manner across participants, despite no shift in central tendency.

To address Hypothesis 1.A., we next focused our analysis on detection of oscillatory peaks in the two sub-bands identified in the analysis of behavioral oscillations of Experiment 1: *low-beta* (14-16 Hz) and *higher-beta* (19-21 Hz). Visual inspection of the results suggested an interaction between task and epoch in the number of *low-beta* vs. *higher-beta* peaks extracted (**Fig. 5B**). A three-way repeated measures ANOVA did not find a significant three-way Beta band × Task × Epoch interaction (*F*(1,15) = 0.8873, *p* = .361). However, a more focused analysis revealed task-specific patterns. In the DSR task, there was a significant crossover pattern whereby the number of *low-beta* and *higher-beta* peaks changed in opposite directions across the delay epochs (*t*(15) = 2.3586, *p* = 0.032), with *higher-beta* decreasing significantly from *Delay 1.1* to *Delay 1.2* (mean change = -0.242 peaks/channel, *t*(15) = -2.41, *p* = 0.029); an effect most prominent at frontal and occipital electrodes (**Supplementary Figure 4**) and *low-beta* showing a numeric increase (mean change = 0.178 peaks/channel, *t*(15) = 2.02, *p* = 0.061); a trend most prominent at occipital and parietal electrodes (**Supplementary Figure 4**). The NEU condition, in contrast, showed stable patterns in these two sub-bands across the two epochs (all *p*-values > 0.6).

##### Aperiodic spectral component

For the aperiodic offset, the ANOVA revealed a significant main effect of Epoch, (*F*(1,15) = 11.07, *p* = .005), with offset increasing from *Delay 1.1* (*M* = 1.19) to *Delay 1.2* (*M* = 1.32). This indicated elevated broadband power during the post-cue portion of the delay period. There was no main effect of Task (*F*(1,15) = 0.23, *p* = .64) and no Task × Epoch interaction (*F*(1,15) = 3.56, *p* = .08; **Fig. 5C**). For the aperiodic exponent, spectral slopes were numerically steeper during *Delay 1.2* versus *Delay 1.1*, (*F*(1,15) = 3.63, *p* = .076), with no effect of Task (*F*(1,15) = 0.58, *p*= .46), and no interaction (*F*(1,15) = 2.39, *p* = .14). Non-parametric tests collapsing across tasks confirmed significant epoch effects for both offset (Wilcoxon signed-rank, *z* = -2.79, *p* = .005, *r* = .70) and exponent (*z* = -2.22, *p* = .026, *r* = .56).

##### Burst analysis

(Test of Hypothesis 1.B.) Many analyses of intra- and extracranial recordings have shown that beta-band effects observed in trial-averaged electrophysiological data can be shown to be due to bursting—rather than to a sustained oscillating signal—at the single-trial level. Because the EEG effects relating to the behavioral oscillations from Experiment 1 (**Fig. 5**) emerged from a spectral parameterization routine that is carried out on trial-averaged power spectra, it was important that we assess whether these effects might also be due to bursting in the *low-beta* and *higher-beta* frequency bands.

We carried out our burst analyses with routines that had previously been applied to EEG data from a working memory task to demonstrate that averaged power dynamics in the traditionally defined beta band (15-29 Hz) could be accounted for by burst dynamics observed at the single-trial level (Kavanaugh et al., 2026). Thus, to validate our approach, we first applied a burst analysis to the 4 sec epochs of *Delay 1.2* from the NEU task, from the P3 electrode, bandpass filtered to 15-29 Hz (“broadband beta”). This yielded burst frequencies of 0.91 bursts/sec (SD = .13) for NEU; and of 0.89 bursts/sec (SD = .13) for DSR. These values are similar to the 1.21 bursts/sec (SD = .22) obtained by (Kavanaugh et al., 2026; B.C. Kavanaugh, personal communication). In the narrower *low-beta* (14-16 Hz) and *higher-beta* (19-21 Hz) bands, however, bursts were sparse events that occurred infrequently at any individual electrode (3-7% of trials per channel), although when considering all channels simultaneously, in *low-beta*, at least one burst occurred on 49.5-65.3% of trials (depending on delay epoch and condition) and in *higher-beta*, on 66.9-76.3% of trials. Despite the rare occurrence of bursts in these two frequency bands, the burst rate in both showed a significant increase from *Delay 1.1* to *Delay 1.2*, albeit with no differences between the DSR and NEU tasks, with values for *Delay 1.1* ranging from 0.086-0.13 bursts/sec and values for Delay 1.2 ranging from 0.139-0.176 events/sec (**Fig. 5D**). Separate repeated-measures ANOVAs (Task × Epoch) for each frequency band confirmed significant main effects of Epoch (low-beta: *F*(1,15) = 19.72, *p* < 0.001; higher-beta: *F*(1,15) = 16.66, *p* < 0.001), with no main effects of Task (both *p*-values > 0.38) and no Task × Epoch interactions (both *p*-values > 0.29). Within-task paired *t*-tests confirmed significant increases from *Delay 1.1* to *Delay 1.2* for both bands in both tasks (all *p* ≤ 0.004). Comparing burst rates across frequency bands, *higher-beta* bursts were more frequent than *low-beta* bursts during *Delay 1.1* (both tasks: *t*(15) > 5.37, *p* < 0.001), but this difference diminished during *Delay 1.2* (DSR: *t*(15) = 2.25, *p* = 0.04; NEU: *t*(15) = 3.45, *p* = 0.004), reflecting a proportionally larger increase in *low-beta* burst activity in the early delay period following the cues.

##### Power

To characterize the phase-locking properties of beta-band activity, we decomposed raw EEG power in the *low-beta* and *higher-beta* bands into induced (non-phase-locked) and evoked (phase-locked) components. Across both tasks and both frequency bands, power was almost exclusively induced, with evoked power constituting only ∼1-2% of total power (**Supplementary Figure 5**).

Induced power did not mirror the task-specific *Delay 1.1* to *Delay 1.2* crossover pattern observed in the peak extraction analysis (**Supplementary Figure 5A**). In *low-beta*, there was a main effect of Task (*F*(1,15) = 5.79, *p* = .030), with NEU showing greater induced power than DSR across both epochs, but no Task × Epoch interaction (*F*(1,15) = 1.72, *p* = .210). In *higher-beta*, there was a main effect of Epoch (*F*(1,15) = 9.38, *p* = .008), with induced power increasing from *Delay 1.1* to *Delay 1.2* in both tasks, no interaction (*F*(1,15) = 1.48, *p* = .242). The absence of interactions indicates that raw power measures—which conflate periodic oscillations with aperiodic activity—do not capture the subtle, task-specific shifts in oscillatory peak distributions revealed by the spectral parameterization. The epoch effect in *higher-beta* power may reflect the task-general increases in burst rate and aperiodic offset reported above.

Evoked power, which reflects phase-locked oscillatory activity, revealed a different pattern (**Supplementary Figure 5B**). Both frequency bands showed significant main effects of Task (*low-beta*: *F*(1,15) = 7.83, *p* = .014; *higher-beta*: *F*(1,15) = 20.11, *p* < .001) and Epoch (*low-beta*: *F*(1,15) = 9.38, *p* = .008; *higher-beta*: *F*(1,15) = 13.20, *p* = .003), with no interactions (both *p*-values > .37). In both bands, NEU evoked power was greater than DSR, particularly during the post-cue epoch. This is consistent with a more stereotyped, phase-consistent neural response to the uninformative cue in the NEU task, and more variable power timing across trials in response to the retrocue in the DSR task.

Finally, to determine whether task differences reflected differential cue-related modulation rather than baseline differences, we computed percent power change from pre-cue to post-cue using *Delay 1.1* as baseline. Induced power showed no task differences in percent change (*low-beta*: *t*(15) = -1.25, *p* = .23; *higher-beta*: *t*(15) = - 1.22, *p* = .24; **Supplementary Figure 5C**), indicating that both tasks exhibited similar cue-related increases in non-phase-locked power. By contrast, evoked power revealed a significant task difference in *low-beta* (*t*(15) = -3.46, *p* = .004): DSR exhibited a larger decrease in phase-locked power (-48.1 ± 8.2%) compared to NEU (-24.9 ± 7.4%). *Higher-beta* showed a similar numerical pattern but did not reach significance (DSR: - 35.0 ± 9.6%; NEU: -19.7 ± 7.5%; *p* = .17). This pattern is consistent with a more disruptive effect of the informative retrocue (DSR) compared to the uninformative cue (NEU) on cross-trial phase consistency.

#### Interim summary—EEG spectral analysis

Filtering the EEG data into the two narrow bands identified in Experiment 1—*low beta* (14-16 Hz) and *higher-beta* (19-21 Hz)—produced results that are broadly congruent with the behavioral oscillations observed in Experiment 1 (and, therefore, consistent with Hypothesis 1.A). Specifically, in the DSR task, the prioritization cue triggered a shift toward more prominent oscillatory activity in the *low-beta* band (together with a shift away from *higher-beta*). In the NEU task, in contrast, there was only evidence of a shift toward more prominent activity in the alpha band. Analyses of the aperiodic component of the data revealed effects that were common to both tasks. For broadband power (i.e., the offset parameter), there was an increase from *Delay 1.1* to *Delay 1.2*, consistent with a ramping of neural activity leading up to the response. For spectral slope (i.e., the exponent parameter) there was evidence, albeit weaker, for a steeper slope during *Delay 1.2*, which would be consistent with a shift in E:I balance toward greater relative inhibition. Beta-burst rates, although in the expected range within the classically defined beta band, were too sparse in the two narrow bands of theoretical interest to be candidate substrates of the behavioral oscillations observed in Experiment 1 (and, therefore, failed to support Hypothesis 1.B.). Nonetheless, consistent with the oscillatory dynamics, burst rates increased from pre-cue to post-cue periods in both the *low-beta* and *higher-beta* bands, albeit with no task differences. Finally, raw power measures did not capture the task-specific oscillatory signatures identified through spectral parameterization, although evoked power analyses revealed that the informative retrocue in DSR was more disruptive to cross-trial phase consistency than the uninformative cue in NEU.

#### Priority-related transformation of signals (for Hypothesis 2)

##### Representational Similarity

Across the *Delay 1.1* and *Cue 1 epochs*, RDMs computed from 28-sec windows exhibited moderate levels of correlation (ranging from -.0154 to 0.3456; **Fig. 6**). For the cross-epoch temporal generalization RSA, however, correlations were markedly lower (-.0344, to .1282) and returned no significant clusters when quantifying evidence for generalization from *Delay 1.1* to *Cue 1*.

**Fig. 6.**
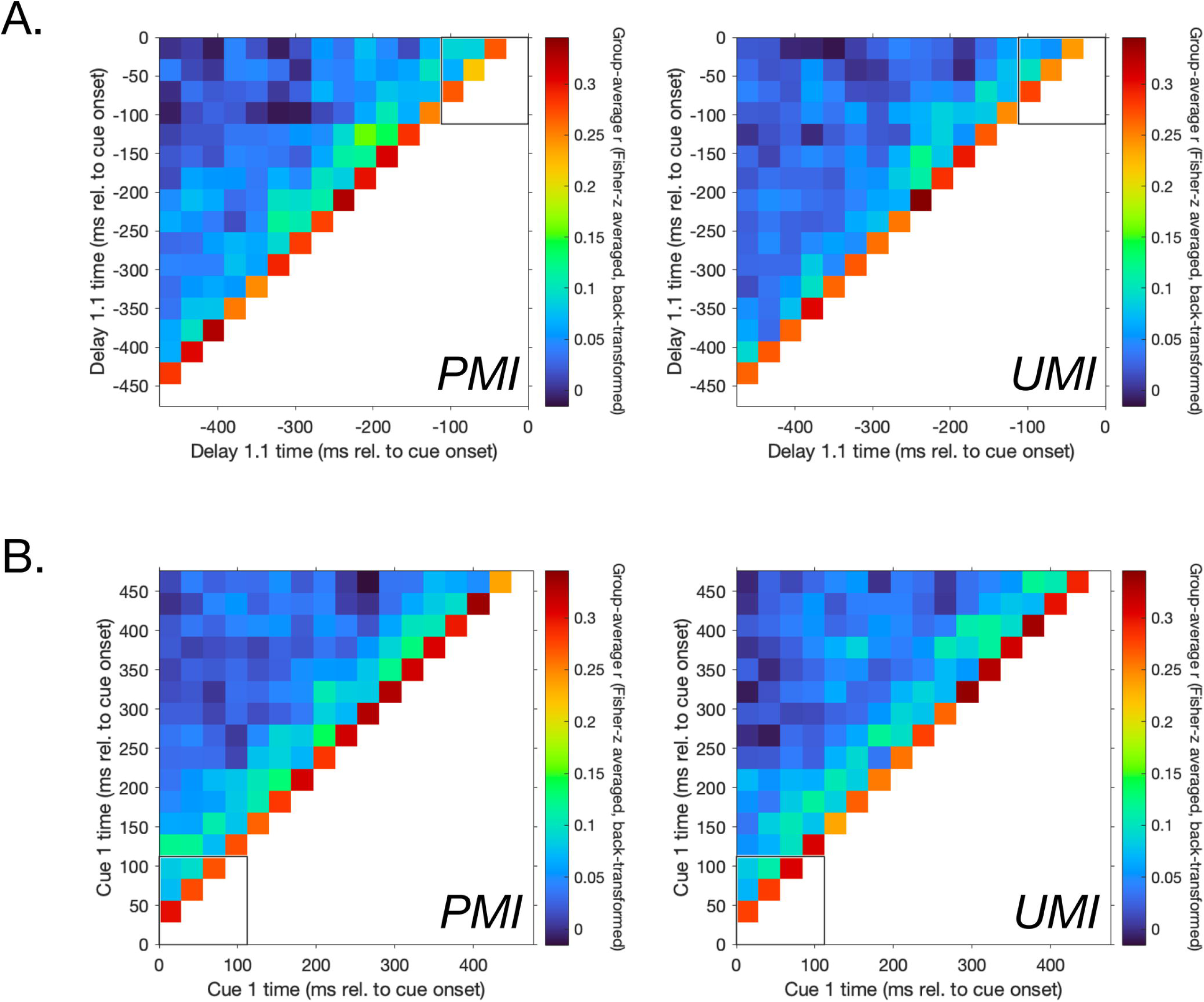
Temporal generalization RSA. **A.** Group-averaged within-epoch self-consistency matrices for the prioritized memory item (PMI, left column) and unprioritized memory item (UMI, right column) during *Delay 1.1.* **B.** Same as **A.** for *Cue 1.* Each matrix shows the Fisher z-averaged, back-transformed correlation between representational dissimilarity matrices (RDMs) computed in successive 28-ms windows within the same epoch, with only the lower triangle displayed (diagonal and upper triangle omitted, as these reflect trivial self-comparisons and their mirror-symmetric duplicates, respectively). Axes indicate window time in milliseconds relative to epoch onset. The outlined square in each panel marks the 112-ms window immediately adjacent to the cue boundary, corresponding to the near-cue self-consistency analysis reported separately in the text.

For the *Delay 1.1* temporal generalization RSA (**Fig. 6A**), reliable self-consistency was observed for both memory items within the full 500-ms epoch (future PMI: *r* = .077, *p* < .001; future UMI: *r* = .069, *p* < .001), as well as within the 112-ms window immediately preceding the cue (future PMI *r* = .167, future UMI *r* = .165, both *p* < .001). Similar results were observed for the *Cue 1* temporal generalization RSA (**Fig. 6B**), for the full 500-ms window (PMI: *r* = .079, *p* < .001; UMI: *r* = .079, *p* < .001) and for the 112-ms window beginning with cue onset (PMI *r* = .184, UMI *r* = .190, both *p* < .001). No PMI–UMI difference in self-consistency was observed in either epoch for either window size (all *p* ≥ .246). Furthermore, self-consistency did not differ significantly between *Delay 1.1* and *Cue 1*, for either item at either window size (500-ms: PMI: *t*(15) = -0.18, *p* = .857, UMI: *t*(15) = -1.49, *p* = .157; 112-ms: PMI: *t*(15) = -0.98, *p* = .345, UMI: *t*(15) = -1.54, *p* = .144). This indicated that representational similarity within the two epochs was comparably self-consistent, ruling out the possibility that *Cue 1* was simply a degraded or noisier version of patterns in *Delay 1.1*.

For the window that spanned cue onset (i.e., incorporating the end of *Delay 1.1* and the beginning of *Cue 1*), self-consistency was statistically significant (PMI: *t*(15) = 8.34, *p* < .001, UMI: *t*(15) = 11.05, *p* < .001), but also lower, for both the PMI and the UMI, in comparison to the pooled within-epoch estimate across *Delay 1.1* and *Cue 1* (unmatched lags: PMI: within *r* = .176 vs. cross *r* = .083, UMI within *r* = .177 vs. cross *r* = .076, both *p*-values < .0001; matched lags restricted analysis: PMI within *r* = .176 vs. cross *r* = .129, *p* = .0051; UMI within *r* = .177 vs. cross *r* = .113, *p* < .0001). Finally, these drops in self-consistency for the cue-onset-spanning window did not differ in magnitude between the PMI and UMI (*p* = .305).

##### Self-correlation of EEG topographies

Self-correlation of the raw EEG from the DSR task, for both the PMI and UMI, exhibited large drops when comparing *Delay 1.1* to *Cue 1* (PMI: observed *r* = 0.299, ceiling *r* = 0.847, *t*(15) = 14.83, *p* < .0001; UMI: observed *r* = 0.334, ceiling *r* = 0.915, *t*(15) = 15.34, *p* < .0001; **Fig. 7**), indicating substantial priority-related reorganization for both stimuli. Furthermore, the magnitude of this drop differed between the two, with that of the UMI (1.244 Fisher-z) significantly greater than that of the PMI (0.971 Fisher-z; paired-*t*(15) = −3.46, *p* = .0035). This result was robust under leave-one-out resampling with every one of the 16 single-participant exclusions preserving significance (with *t*-values ranging from −3.10 to −3.91 and *p*-values ranging from .0016 to .0079). For the NEU task, although cue-related drops in self-correlation for tested and untested items were significant (both *p*-values ≤ .0012; **Fig. 7**), these were significantly smaller than those for the DSR task (both *p*-values < .001), and the magnitude of the drop did not differ between them (*t*(14) = −0.07, *p* = .945). Therefore, these results are consistent with Hypothesis 2.

**Fig 7.**
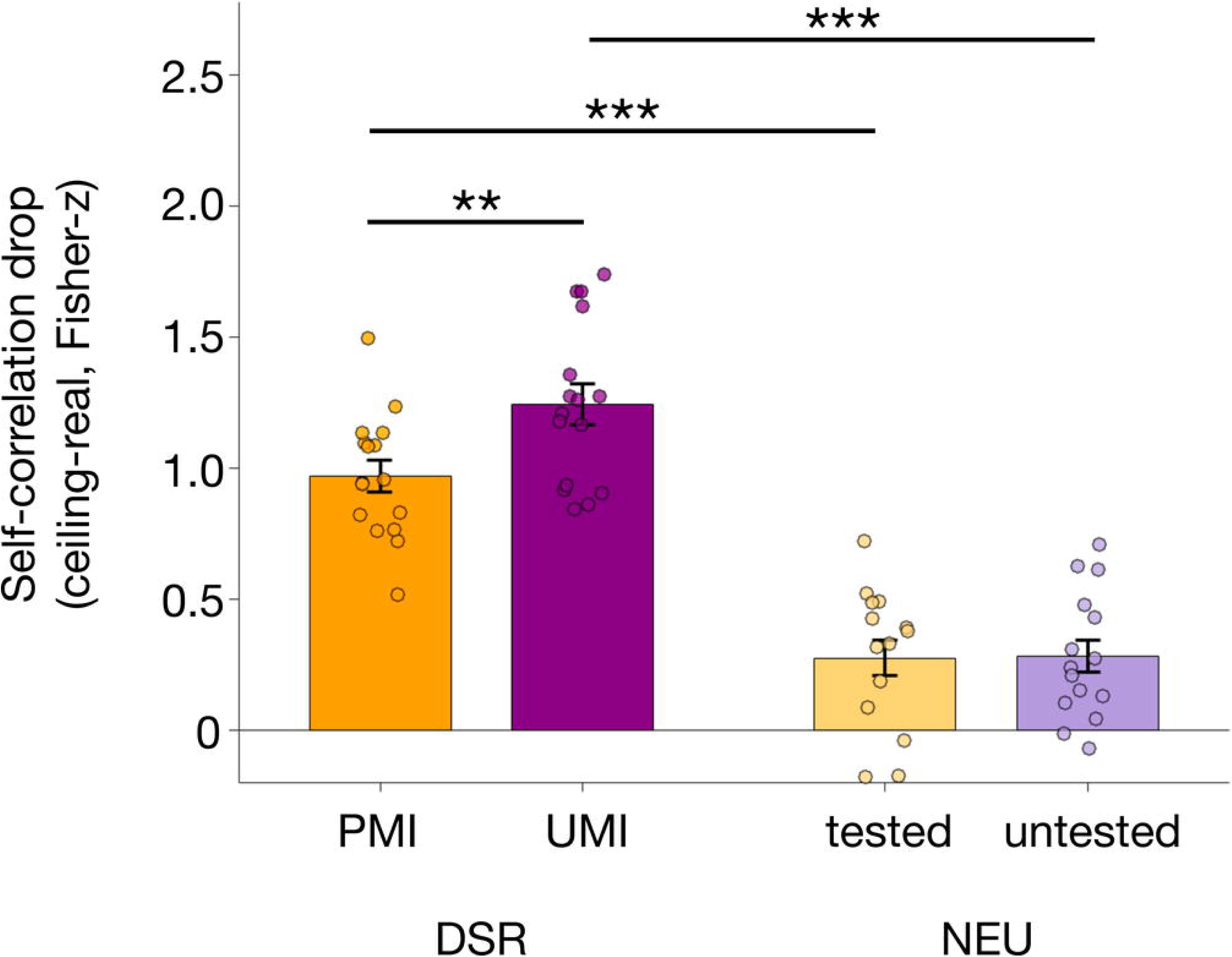
EEG cue-locked self-correlation drop. Self-correlation drop was defined as the bootstrapped noise ceiling (the correlation expected from trial-sampling noise alone, estimated from two independent resamples of data from the *Delay 1.1* epoch) minus the observed (‘real’) *Delay 1.1*-to-*Cue 1* correlation, in Fisher-z units, per participant. Bars show group mean ± 1 *SEM* for the two memory items of each task (DSR-PMI, DSR-UMI, NEU-tested, NEU-untested); individual points show participant-level values, jittered horizontally for visibility. Brackets indicate significant paired comparisons. ** corresponds to *p* < .01; *** corresponds to *p* < .001.

#### Interim discussion—Experiment 2

Behavioral results broadly replicated the evidence from Experiment 1 that prioritization shields an item from the biasing influence of other information recently or currently in working memory. One possible exception to this was the fact that, unique to Experiment 2, recall of *Sample 2* showed greater repulsion from *Sample 1* in DSR than in NEU. However, we speculate that this, together with other specific differences between the two experiments, may reflect an interaction of prioritization with the longer delay period of Experiment 2. For example, previous work has suggested that longer delays (Chunharas et al., 2022), as well as “active, not passive, maintenance” (Scotti et al., 2021) are associated with greater separation between items.

Consistent with Hypothesis 1.A, spectral parameterization of the EEG in the DSR task revealed a cue-locked increase of oscillatory peaks in the *low-beta* band, together with a decrease in oscillatory peaks in the *higher-beta* band, suggesting an underlying eletrophysiological basis for the priority-specific behavioral oscillations observed in Experiment 1. Inconsistent with Hypothesis 1.B, however, burst analyses indicated that these effects in these narrow frequency bands were unlikely to be due to changes of burst dynamics at the single trial level.

With regard to Hypothesis 2, the larger drop in EEG self-correlation for the UMI than for the PMI is consistent with the prediction that cue-locked priority-based transformations would be more pronounced for the uncued item (i.e., the UMI) than for the cued item (the PMI). However, this conclusion needs to be qualified by the fact that this analysis did not measure representation directly.

## Discussion

The results from these two experiments provide evidence for a model in which the prioritization of one among two items held in working memory prompts representational transformations that may be larger for the uncued item (i.e., the UMI) than for the cued item (the PMI), and in which the encoding of priority status is implemented, at least in part, by dynamics in the beta band. Previous studies of the DSR task combining EEG and transcranial magnetic stimulation (TMS) have shown that decoding of the UMI depends on patterns in the beta band (Rose et al., 2016; Fulvio & Postle, 2026), and this effect is associated with cue- and TMS-related dynamics in the low beta band (i.e., 13-20 Hz; Fulvio & Postle, 2026). Furthermore, single pulses of TMS (spTMS) delivered during *Delay 1.2* can produce behavioral evidence for the involuntary retrieval of the UMI (indexed by an inflated false-alarm rate; Rose et al., 2016; Fulvio & Postle, 2020), and the magnitude of this behavioral effect is predicted by the magnitude of a spTMS-related decrease in beta-band components derived from a decomposition of the EEG (Fulvio et al., 2024). In the present study, we have produced evidence that prioritization is associated with a behavioral oscillation in the low-beta band and by an increase of oscillatory peaks in this same narrow frequency band in the EEG. Thus, our working model is that UMI status is encoded, and sustained, by a process that cycles at a frequency in the low-beta band. Although this account lacks an explicit mechanism, future work can explore the possibility that this may be due to the encoding of the two memory items at different phases of a low-beta oscillation (Abdalaziz et al., 2023).

Although we do not yet understand the precise mechanism that implements the state of deprioritization, we speculate that its dependence on oscillations at a low-beta frequency links it to the natural frequency of thalamocortical circuits at posterior nodes of the frontoparietal priority map. This follows from the results of several studies that have recorded the EEG while spTMS is delivered to posterior parietal cortex (PPC): spTMS of PPC produces a spectral perturbation that is strongest in β1 (13-20 Hz; in comparison to spTMS of occipital cortex—alpha: 8-12 Hz—and spTMS of premotor cortex--β2: 21-29 Hz; Rosanova et al., 2009); when measured as the global mean field amplitude (GMFA), the amplitude of the response evoked by spTMS of PPC varies with pre-spTMS phase in the beta band (Kundu et al., 2014); and spTMS-evoked phosphene perception depends on pre-spTMS power in the beta band for spTMS of PPC (in comparison to its dependence on pre-spTMS power in the alpha band for spTMS of occipital cortex; Samaha et al., 2017). Assuming that priority in working memory is encoded in the frontoparietal priority map, it may be that the top-down signals that impose this priority status on stimulus representations in early visual areas oscillate at the natural frequency of the PPC.

The behavioral and EEG data presented here point to an oscillatory mechanism supporting prioritization: recall precision oscillates at 15 Hz, and the prioritization-specific increase of oscillatory peaks in this narrow frequency band cannot be accounted for by the sparse bursting rate that we observed. This suggests that prioritization is supported by a mechanism that is fundamentally different from the beta bursts, also observed at posterior electrodes, that implement inhibitory cognitive control (e.g., Lundqvist et al., 2024; Wessel & Anderson, 2024). Importantly, however, our claim does not extend to the processes that underlie the more general function of the maintenance of information in working memory. Indeed, when we applied a burst analysis to the broader band of traditionally defined beta (15-29 Hz), we observed a burst rate comparable to one that has been shown to underlie load-related dynamics in the EEG (Kavanaugh et al., 2026).

Several analyses suggest that there are cue-locked transformations for both memory items. The significant cue-related drops in self-consistency of cross-temporal RSA that were observed for both the UMI and PMI were comparable between the two. This would be consistent with the possibility that the transformations of the PMI and UMI are triggered by a common factor, as would occur with the change of a control variable in a dynamical system. Also consistent with this possibility is the fact that trial-by-trial variation in the efficacy of the geometric transformation of the PMI and of the UMI are highly correlated, as measured with fMRI and with EEG (Wan et al., 2024). Despite these considerations, the self-correlation analysis of the raw EEG indicated that the priority-related transformation of the UMI may be larger than that of the PMI, a possibility that would be consistent with previous fMRI studies (e.g., Yu, Teng, & Postle, 2020; Teng & Postle, 2024). We speculate that this may reflect a rotation of the UMI into an action-null subspace, whereas the PMI occupies an action-potent subspace that is geometrically more similar to the representational state of both memory items during portions of the trial when both are equally likely to be needed for the recall response.

In conclusion, the results we report here provide novel insights about why and how prioritization influences working memory performance. Prioritization has the effect of shielding the cued item from the biasing effects of other information currently and previously held in working memory. One consequence of this is a reversal of a commonly observed serial position effect. Behavioral and EEG data provide converging evidence that this shielding effect is implemented, in part, by a mechanism oscillating in the low-beta frequency range. Furthermore, correlation of the raw EEG channel patterns indicates that the effects of this mechanism are most pronounced on the UMI, offering a possible explanation for previous reports of the selective effects of prioritization on patterns of decoding neural representations of the UMI. The oscillatory nature of this mechanism distinguishes it from top-down inhibitory control that is implemented by bursting in the beta band, and we speculate that its frequency profile links it to the operation of a frontoparietal priority map.

## Acknowledgements

We thank Miral Abdalaziz for input and feedback regarding the experimental design. We also thank Dr. Brian Kavanaugh and Dr. Stephanie Jones for their insights about beta burst analyses and sharing of results to validate our findings.

