## Supplementary Figure 1 for "Prioritization in working memory reduces interference via beta band-linked stimulus transformations"

**Supplementary Information**

*Experiment 1 Supplemental analyses*

*Mean absolute error.* Splitting the DSR *Recall 2* responses according to whether the trial was a stay trial (same item cued for both recalls) or a switch trial (different items cued), revealed no difference in performance (**Supplementary** **Fig. 1**).

*Current trial inter-item bias.* When modeling the bias exerted by the current trial’s untested item on recall in isolation, we found, in contrast to MAE, that inter-item bias was significantly larger for NEU recall responses. Specifically, NEU recall responses were significantly more attracted to the untested item in the current trial’s memory set than were both DSR *Recall 1* responses (*t*(15) = -3.558, *p* = .003; **Supplementary** **Fig. 1C**) and DSR *Recall 2* responses (*t*(15) = -2.982 *p* = .009). Separate *t*-tests comparing inter-item bias for NEU recall responses with DSR *Recall 2* on stay trials and switch trials found that the larger bias in NEU responses was significant compared to switch trials (*t*(15) = -3.228, *p =* .0056) but did not reach significance when compared to stay trials (*p =* .149). An LME model also revealed a significant current-trial inter-item angular difference x task interaction (*F*(1,140) = 4.2091, *p* = .042), and a main effect of task on inter-item bias (*F*(1,140) = 9.1491, *p* = .003); the main effect of current trial inter-item angular difference did not reach significance (*p* = .068).

*Previous trial inter-item bias.* We separately considered that prioritization may also impact the bias exerted by previous trial memory information. We carried out traditional serial dependence analyses (see ***Serial dependence*** section in Methods). These analyses confirmed the attractive nature of the bias exerted by the previous trial’s tested memory item in the NEU task observed in the full linear mixed effects model reported in Results (*a* = 0.667 deg, *p* < .001), and a non-significant repulsive bias of the previous trial’s untested item (*a =* -.0965 deg; *p* = .293), with a significant difference between these effects (𝛥*a* = 0.7639*, p <* .01; **Supplementary** **Fig. 1D**). In the DSR task, these analyses revealed a significant attractive bias toward the previous trial’s PMI at *Recall 2* (*a* = 0.659 deg, *p* < .01), which was significantly greater than the significantly repulsive bias of the previous trial’s IMI (*a* = -.66 deg, *p* < .01; 𝛥*a* = 1.321, *p* < .01).

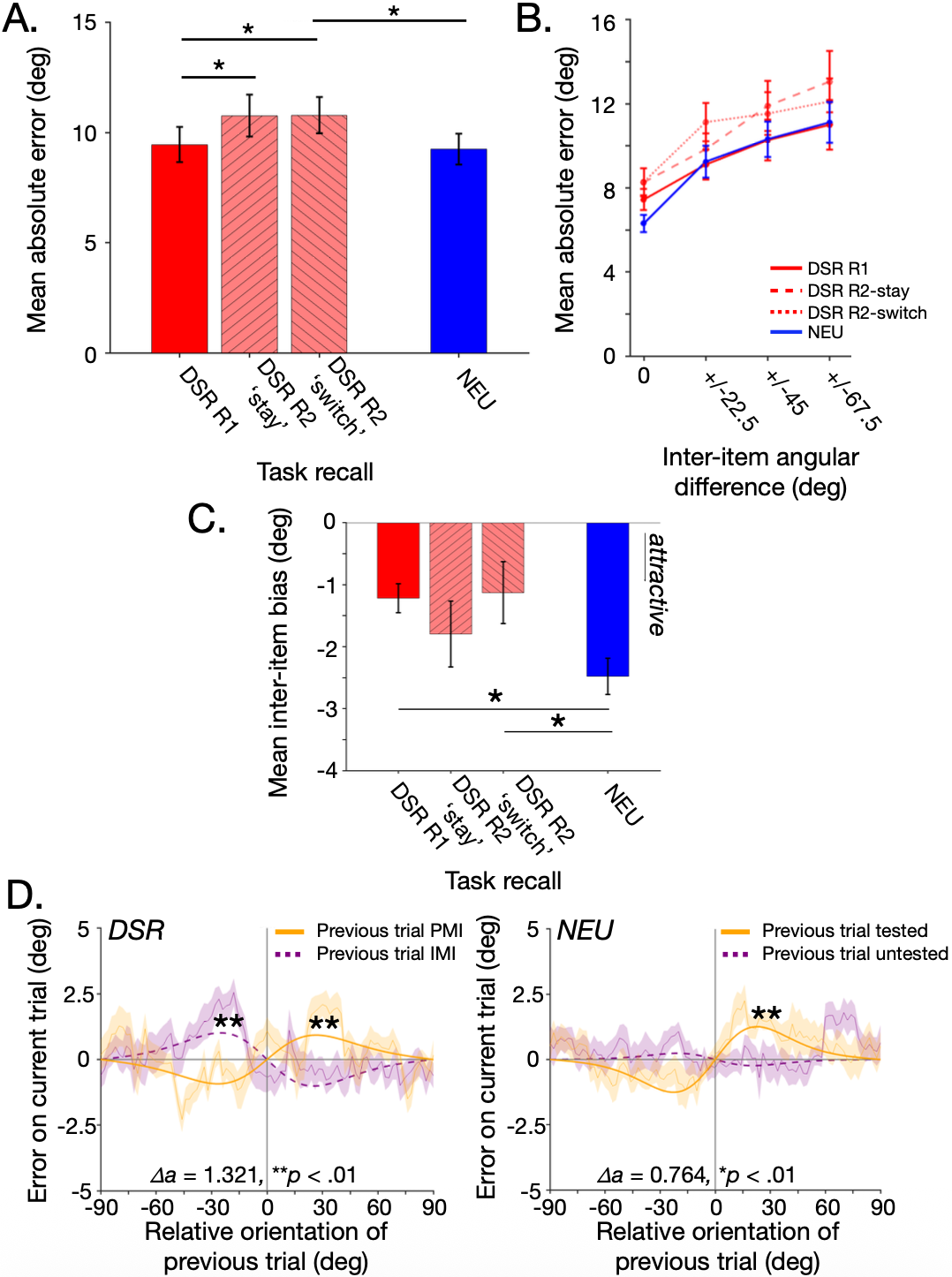

**Figure 1. Experiment 1 recall accuracy and inter-item biases. A.** Mean absolute recall error for the two responses of the DSR task with the second recall accuracy reported separately for ‘stay’ and ‘switch’ trials and the single response of the NEU task. **B.** Same as **A.** separated by the current trial’s inter-item angular difference. See **Figure 2A & 2B** in the main text for the results collapsed over DSR *Recall 2* trial types. **C.** Mean inter-item bias for the two responses of the DSR task with the second recall inter-item bias reported separately for ‘stay’ and ‘switch’ trials and the single response of the NEU task. Negative values indicate that the recall was pulled in the direction of the current trial’s untested item. For panels **A**-**C**, Error bars correspond to +/- 1 SEM. * symbols correspond to significant differences between conditions at the Bonferroni-corrected alpha level = .0083 for 6 paired-sample t-tests. **D.** Group-averaged serial dependence curves and dVM fits to current trial recall error data for the DSR task (left) and the NEU task (right) sorted according to the relative orientation of the previous trial’s PMI (for DSR *Recall 2*)/tested item (orange) and the previous trial’s IMI/untested item (purple). Shaded bands represent ±1 *SEM*. **Bootstrapped *p* < 0.001; 𝛥*a* refers to the diﬀerence in the fitted amplitude parameter between the PMI & IMI (DSR) and tested & untested (NEU) with the corresponding *p*-values derived from permutation testing.

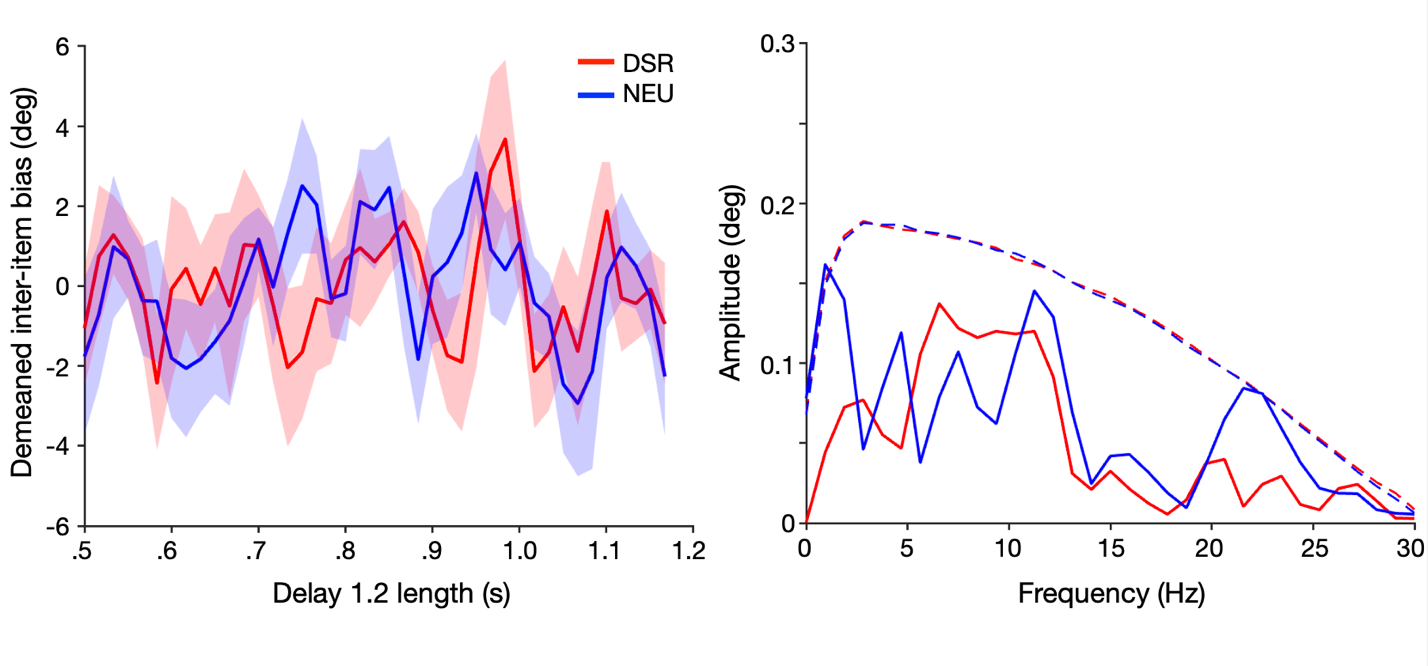

**Figure 2. Experiment 1 behavioral oscillations of mean inter-item bias.** Left: Demeaned mean inter-item bias for DSR *Recall 1* (red) and the single response of the NEU task (blue) as a function of Delay 1.2 length. Error bands correspond to +/- 1 *SEM.* Right: Fast Fourier Transform (FFT) amplitude spectra of mean inter-item bias across delay periods for the DSR (red) and NEU (blue) tasks. Solid lines represent group-averaged amplitudes (in degrees of error). Dashed lines show statistical significance thresholds (95^th^ percentile) derived from permutation testing.

*Experiment 2 Supplemental analyses*

*Mean absolute error.* As in Experiment 1, splitting the DSR *Recall 2* responses according to whether the trial was a stay trial (same item cued for both recalls) or a switch trial (different items cued), revealed no difference in performance (**Supplementary** **Fig. 3**).

*Current trial inter-item bias.* When modeling the bias exerted by the current trial’s untested item on recall in isolation, we found that inter-item bias was significantly larger for NEU recall responses. Specifically, NEU recall responses were significantly more attracted to the untested item in the current trial’s memory set than were both DSR Recall 1 responses (*t*(15) = 4.0410, *p* = .001; **Supplementary** **Fig. 3C**) and DSR Recall 2 responses (*t*(15) = 4.3110, p < .001). Separate *t*-tests comparing inter-item bias for NEU recall responses with DSR *Recall 2* on stay trials and switch trials found that the larger bias in NEU responses was significant compared to switch trials (*t*(15) = -3.332, *p =* .0045) but did not reach significance at the Bonferroni-correct alpha-level of .0083 when compared to stay trials (*p =* .0156). An LME model also revealed a significant current-trial inter-item angular difference x task interaction (F(1,140) = 5.325, p = .022), as well as a main effect of current trial inter-item angular difference (F(1,140) = 11.631, p < .001) and a main effect of task on inter-item bias (F(1,140) = 12.742, p < .001).

*Previous trial inter-item bias.* In the NEU task, there was a significant attractive bias of the previous trial’s tested item (*a =* 1.904 deg, *p* < .001), no significant bias exerted by the previous trial’s untested item (*a =* 0.047 deg, *p* = .539), and the difference between them was significant (𝛥*a* = 1.857*, p <* .01; **Supplementary** **Fig. 3D**). In the DSR task, there was a small but significant attractive bias of the previous trial’s tested item (*a =* 0.151 deg, *p* = .043), and a non-significant repulsive bias exerted by the previous trial’s untested item (*a =* -0.4513 deg, *p* = .386), and the difference between them was not significant (𝛥*a* = 0.6023*, p =* .074).

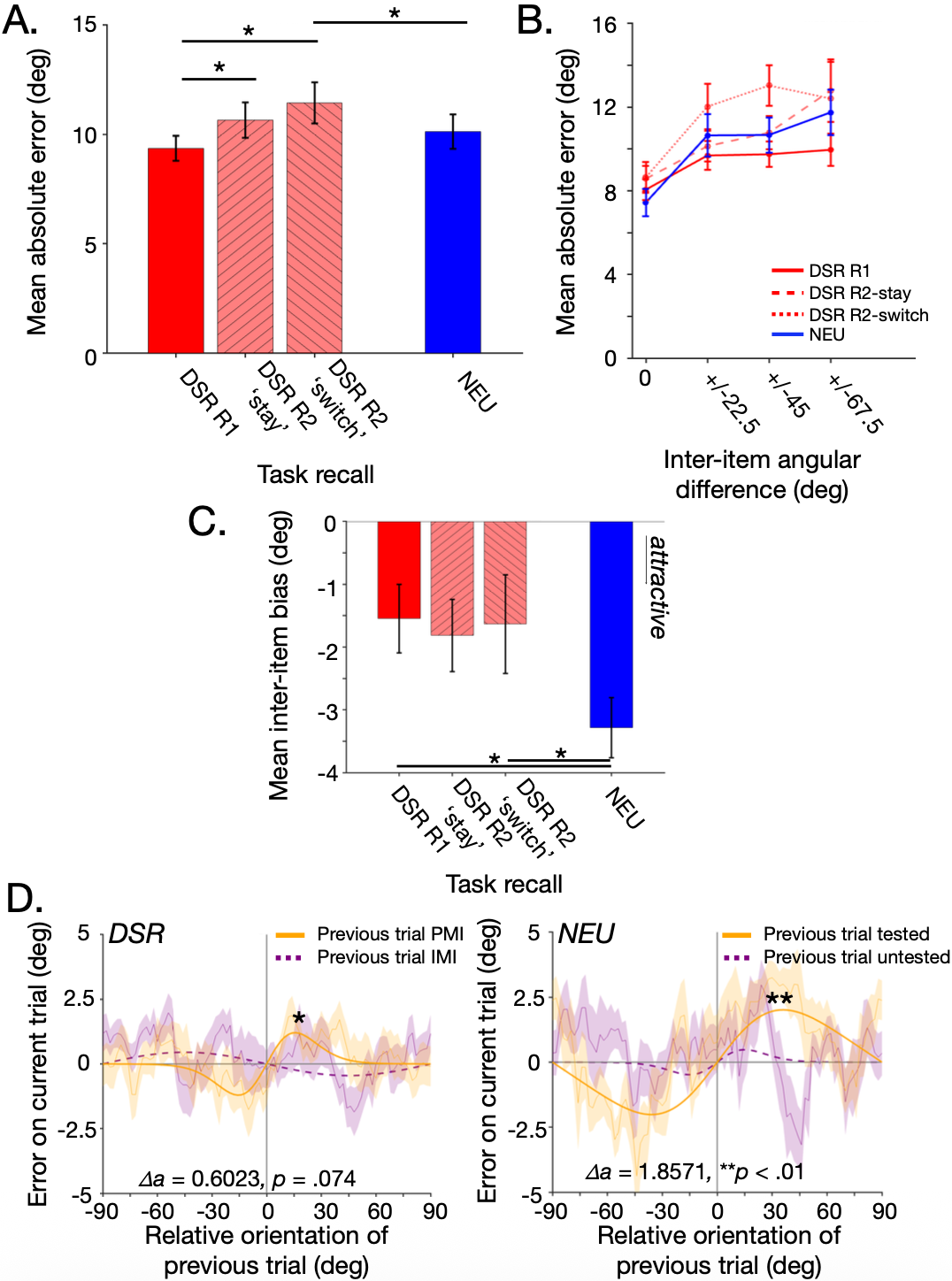

**Figure 3. Experiment 2 recall accuracy and inter-item biases. A.** Mean absolute recall error for the two responses of the DSR task with the second recall accuracy reported separately for ‘stay’ and ‘switch’ trials and the single response of the NEU task. **B.** Same as **A.** separated by the current trial’s inter-item angular difference. See **Figure 4A & 4B** in the main text for the results collapsed over DSR *Recall 2* trial types. **C.** Mean inter-item bias for the two responses of the DSR task with the second recall inter-item bias reported separately for ‘stay’ and ‘switch’ trials and the single response of the NEU task. Negative values indicate that the recall was pulled in the direction of the current trial’s untested item. For panels **A**-**C**, Error bars correspond to +/- 1 SEM. * symbols correspond to significant differences between conditions at the Bonferroni-corrected alpha level = .0083 for 6 paired-sample t-tests. **D.** Group-averaged serial dependence curves and dVM fits to current trial recall error data for the DSR task (left) and the NEU task (right) sorted according to the relative orientation of the previous trial’s PMI (for DSR *Recall 2*)/tested item (orange) and the previous trial’s IMI/untested item (purple). Shaded bands represent ±1 *SEM*. **Bootstrapped *p* < 0.001; 𝛥*a* refers to the diﬀerence in the fitted amplitude parameter between the PMI & IMI (DSR) and tested & untested (NEU) with the corresponding *p*-values derived from permutation testing. Compare with **Supplementary Figure 1.**

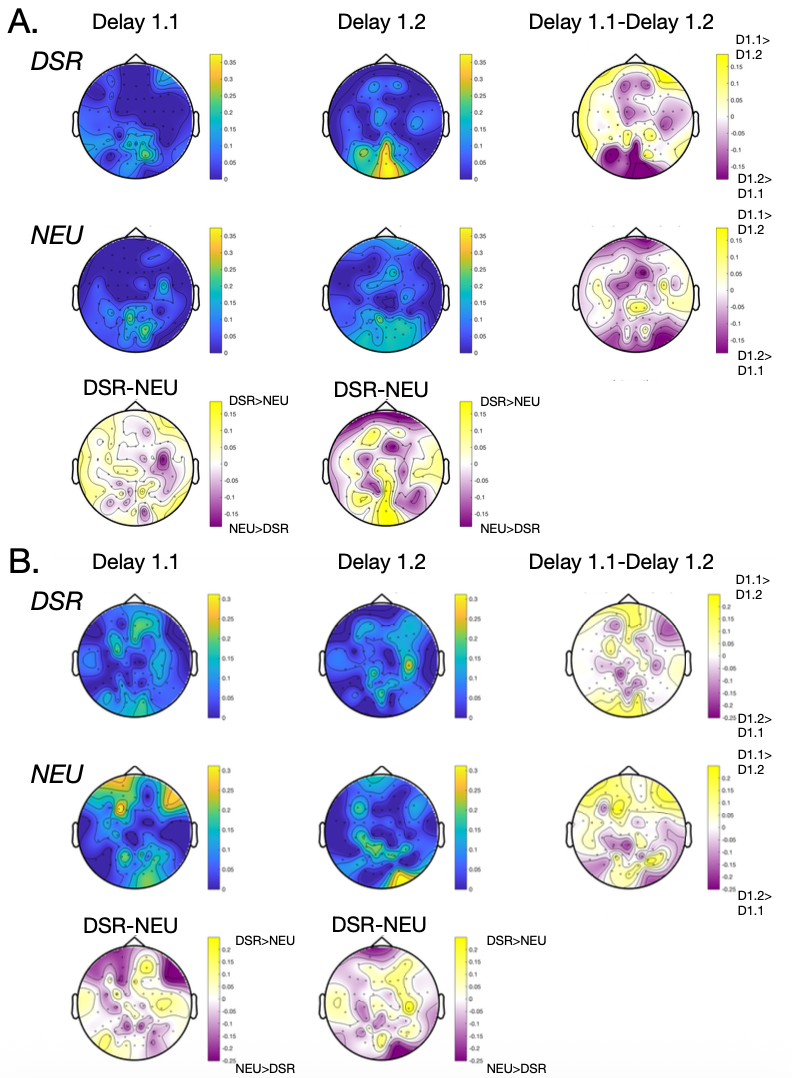

**Figure 4. Topography of beta peak oscillations. A.** *Low-beta* (14-16 Hz) oscillation topographies. Left column: Mean number of extracted peaks in the low beta (14-16 Hz) band per channel for *Delay 1.1* in the DSR task (top row), the NEU task (middle row), and their difference (DSR-NEU, bottom row). Middle column: Same as the left-hand column, but for *Delay 1.2.* Right column: Within-task differences between the two delay periods (*Delay 1.1-Delay 1.*2) for the DSR task (top row) and the NEU task (middle row). **B.** Same as **A.** for *higher-beta* (19-21 Hz) oscillation topographies.

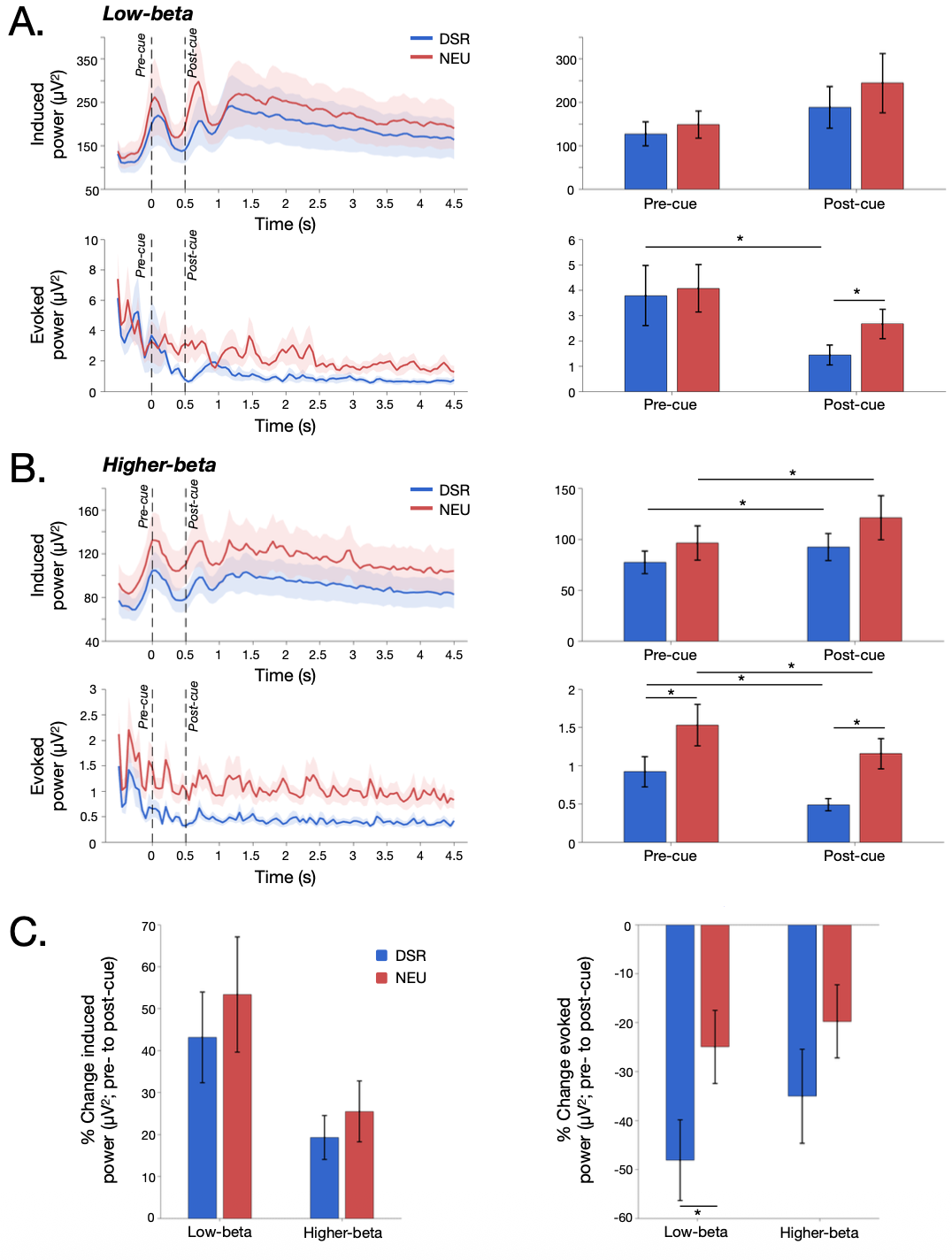

**Figure 5. EEG power analyses. A.** Left: Raw, uncorrected time-resolved induced (upper row) and evoked (lower row) *low-beta* (14-16 Hz) power for the DSR task (blue) and NEU task (red) for the timepoints spanning *Delay 1.1* (‘pre-cue’) through *Delay 1.2* (‘post-cue’). Right: Mean power for the two tasks during the final 400 ms of the *Delay 1.1* and the first 400 ms of *Delay 1.2* (beginning 100 ms after cue offset). **B.** Same as **A.** for *higher-beta* (19-21 Hz) power. **C.** Percent change (post- - pre-cue)/pre-cue *100 in induced power (left) and evoked power (right) for the two tasks and frequency bands. Error bars correspond to +/- 1 *SEM.* * corresponds to significant differences at the Bonferroni-corrected alpha-level = .025.
